# Glycan processing in the Golgi – optimal information coding and constraints on cisternal number and enzyme specificity

**DOI:** 10.1101/2020.05.18.101444

**Authors:** Alkesh Yadav, Quentin Vagne, Pierre Sens, Garud Iyengar, Madan Rao

## Abstract

Many proteins that undergo sequential enzymatic modification in the Golgi cisternae are displayed at the plasma membrane as cell identity markers. The modified proteins, called glycans, represent a molecular code. The fidelity of this *glycan code* is measured by how accurately the glycan synthesis machinery realises the desired target glycan distribution for a particular cell type and niche. In this paper, we quantitatively analyse the tradeoffs between the number of cisternae and the number and specificity of enzymes, in order to synthesize a prescribed target glycan distribution of a certain complexity. We find that to synthesize complex distributions, such as those observed in real cells, one needs to have multiple cisternae and precise enzyme partitioning in the Golgi. Additionally, for fixed number of enzymes and cisternae, there is an optimal level of specificity of enzymes that achieves the target distribution with high fidelity. Our results show how the complexity of the target glycan distribution places functional constraints on the Golgi cisternal number and enzyme specificity.

## I. INTRODUCTION

A majority of the proteins synthesized in the endoplasmic reticulum (ER) are transferred to the Golgi cisternae for further chemical modification by glycosylation [1], a process that sequentially and covalently attaches sugar moieties to proteins, catalyzed by a set of enzymatic reactions within the ER and the Golgi cisternae. These enzymes, called glycosyltransferases, are localized in the ER and cis-medial and trans Golgi cisternae in a specific manner [2, 3]. Glycans, the final products of this glycosylation assembly line are delivered to the plasma membrane (PM) conjugated with proteins, whereupon they engage in multiple cellular functions, including immune recognition, cell identity markers, cell-cell adhesion and cell signalling [2–6]. This *glycan code* [7, 8], representing information [9] about the cell, is generated dynamically, following the biochemistry of sequential enzymatic reactions and the biophysics of secretory transport [4, 10, 11].

In this paper, we will focus on the role of glycans as markers of cell identity. For the glycans to play this role, they must inevitably represent a molecular code [4, 7, 11]. While the functional consequences of glycan alterations have been well studied, the glycan code has remained an enigma [7, 11–13]. In this paper, we study one aspect of molecular coding, namely the *fidelity* of this molecular code generation. While it has been recognised that fidelity of the glycan code is necessary for reliable cellular recognition [14], a quantitative measure of fidelity of the code and its implications for cellular structure and organization are lacking.

There are two aspects of the cell-type specific glycan code that have an important bearing on quantifying fidelity. The first is that extant glycan distributions have high *complexity*, owing to evolutionary pressures arising from (a) reliable cell type identification amongst a large set of different cell types in a complex organism, the preservation and diversification of “self-recognition” [5], (b) pathogen-mediated selection pressures [2, 4, 6], and (c) *herd immunity* within a heterogenous population of cells of a community [15] or within a single organism [5]. Here, we will interpret this to mean that the *target distribution* of glycans of a given cell type is complex; in Sect. II we define a quantitative measure for complexity and demonstrate its implications in the context of *human* T-cells. The second is that the cellular machinery for the synthesis of glycans, which involves sequential chemical processing via cisternal resident enzymes and cisternal transport, is subject to variation and noise [4, 10, 11]; the *synthesized glycan distribution* is, therefore, a function of cellular parameters such as the number and specificity of enzymes, inter-cisternal transfer rates, and number of cisternae. We will discuss an explicit model of the cellular synthesis machinery in Sect. III.

In this paper, we define fidelity as the minimum achievable Kullback-Leibler (KL) divergence [16, 17] between the synthesized distribution of glycans and the target glycan distribution. This KL divergence is a function of the cellular parameters governing glycan synthesis, such as the number and specificity of enzymes, inter-cisternal transfer rates, and number of cisternae (Sect. V). We analyze the tradeoffs between the number of cisternae and the number and specificity of enzymes, in order to achieve a prescribed target glycan distribution with high fidelity (Sect. VI). Our analysis leads to a number of interesting results, of which we list a few here: (i) In order to construct an accurate representation of a complex target distribution, such as those observed in real cells, one needs to have multiple cisternae and precise enzyme partitioning. Low complexity target distributions can be achieved with fewer cisternae. (ii) This definition of fidelity of the glycan code, allows us to provide a quantitative argument for the evolutionary requirement of multiple-compartments. (iii) For fixed number of enzymes and cisternae, there is an optimal level of specificity of enzymes that achieves the complex target distribution with high fidelity. (iv) Keeping the number of enzymes fixed, having low specificity or sloppy enzymes and larger cisternal number could give rise to a diverse repertoire of functional glycans, a strategy used in organisms such as plants and algae.

Stated another way, our results imply that the pressure to achieve the target glycan code for a given cell type, places strong constraints on the cisternal number and enzyme specificity [18]. This would suggest that a description of the nonequilibrium assembly of a fixed number of Golgi cisternae must combine the dynamics of chemical processing with membrane dynamics involving fission, fusion and transport [19, 20], opening up a new direction for future research.

## II. COMPLEXITY OF GLYCAN CODE IN REAL CELLS

Since each cell type (in a niche) is identified with a distinct glycan profile [4, 7, 11], and this glycan profile is noisy because of the stochastic noise associated with the synthesis and transport [11–13], a large number of different cell types can be differentiated only if the cells are able to produce a large set of glycan profiles that are distinguishable in the presence of this noise. A more complex or richer class of glycan profiles is able to support a larger number of well separated profiles, and therefore, a larger number cell types, or equivalently, a more complex organism^1^

In order to implement a quantitative measure of complexity, we first need a consistent way of smoothening or coarse-graining the discrete glycan distribution to remove measurement and synthesis noise. In this paper, we approximate the glycan profile as mixture of Gaussian densities with specified number of components that are supported on a finite set of indices [21]. Since the complexity of *k*-component Gaussian is an increasing function of *k*, we use the number of component *k* and complexity interchangeably.

Using this definition we demonstrate that the glycan profiles of typical mammalian cells are very complex. We obtain target profiles for a given cell type from Mass Spectrometry coupled with determination of molecular structure (MSMS) measurements [22]. Fig. 1 shows the the MSMS data from *human* T-cells and *human* and *mouse* neutrophils [22], and their coarse-grained representations using Gaussian mixture models (GMM) of differing complexity - a low complexity *k* = 3 GMM and high complexity *k* = 20 GMM. It is clear from Fig. 1, that the more complex *k* = 20 GMM is a better representation of the MSMS data as compared to the less complex *k* = 3 GMM. Indeed the *k* = 20 Gaussian mixture model is the best compromise between faithfulness of the representation and cost of an additional component, as seen from the saturation of the likelihood function [17]. Details of this systematic coarse-graining procedure appear in Sect. VI B and Appendix G.

**FIG. 1.**
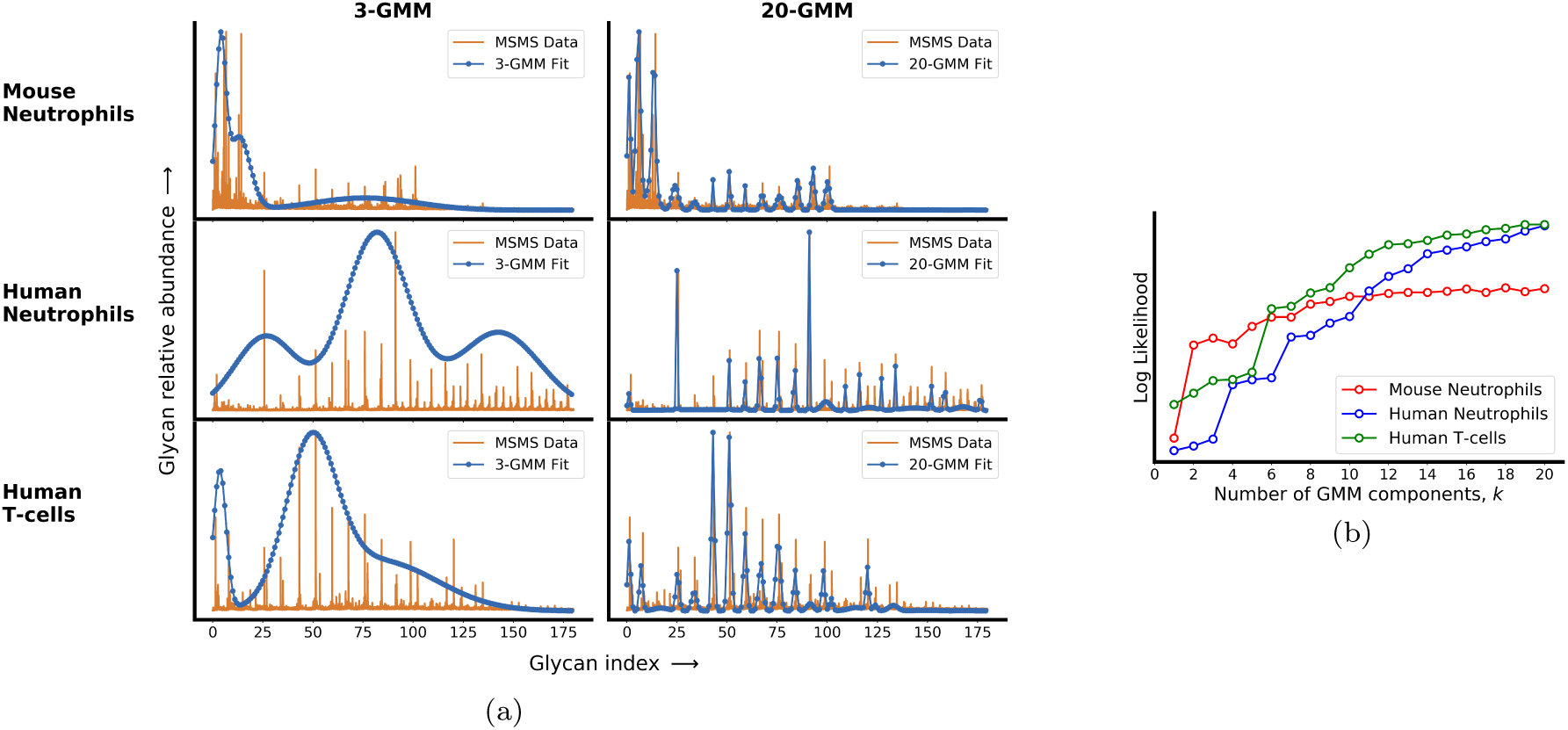
Real cells display a complex glycan distribution. (a) Here we take the MSMS data from *mouse* neutrophils, *human* neutrophils and *human* T-cells and approximate these using Gaussian Mixture Models (GMM) of less complexity 3-GMM (left) and more complexity 20-GMM (right). (b) The change in log likelihood with increase in the number of GMM components for *mouse* and *human* neutrophils and *human* T-cells, shows a saturation at large enough values of *k*, indicating that these glycan distribution are complex. Details appear in Appendix G.

Having demonstrated the complexity of the typical glycan distributions associated with a given cell type, we will now describe a general model of the cellular machinery that is capable of synthesizing glycans of the desired complexity. We expect that cells need a more elaborate mechanism to produce profiles from a more complex set.

## III. SYNTHESIS OF GLYCANS IN THE GOLGI CISTERNAE

The glycan display at the cell surface is a result of proteins that flux through and undergo sequential chemical modification in the secretory pathway, comprising an array of Golgi cisternae situated between the ER and the PM, as depicted in Fig. 2. Glycan-binding proteins (GBPs) are delivered from the ER to the first cisterna, whereupon they are processed by the resident enzymes in a sequence of steps that constitute the N-glycosylation process [2]. A generic enzymatic reaction in the cisterna involves the catalysis of a group transfer reaction in which the monosaccharide moiety of a simple sugar donor substrate, e.g. UDP-Gal, is transferred to the acceptor substrate, by a Michaelis-Menten (MM) type reaction [2]

**FIG. 2.**
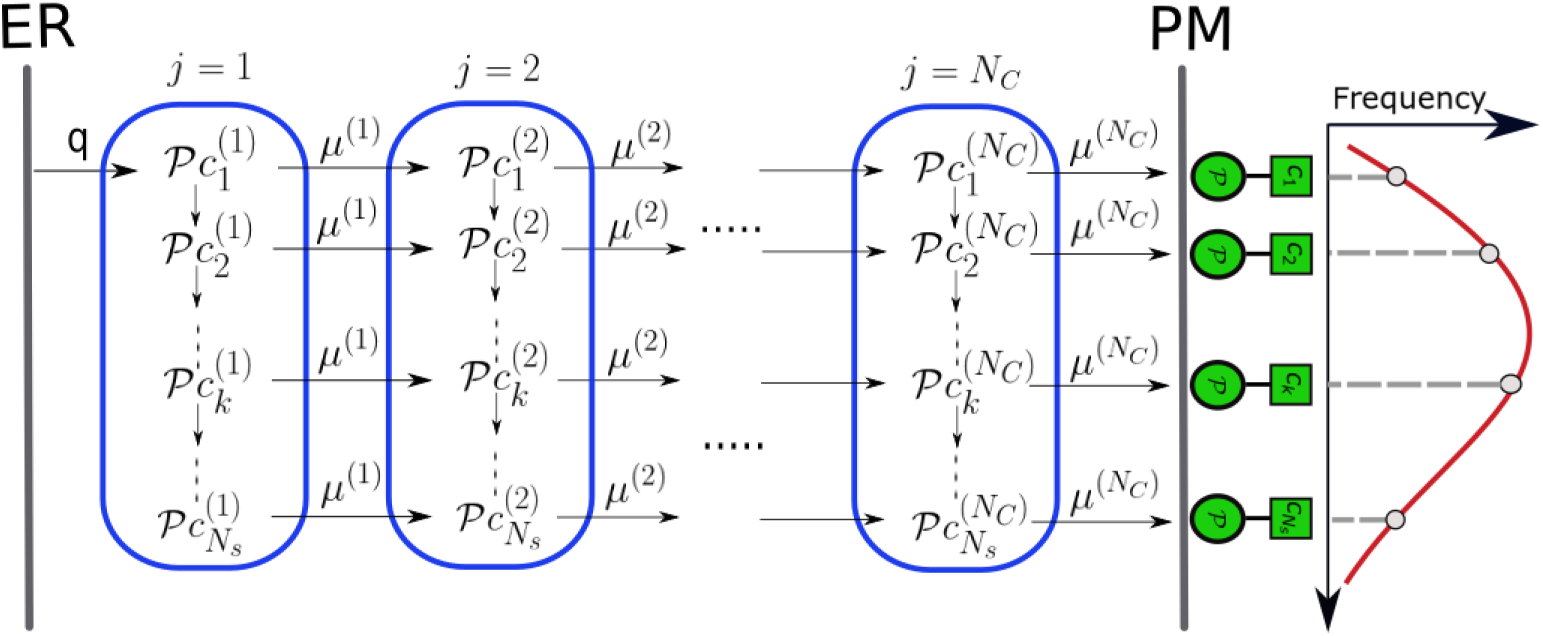
Enzymatic reaction and transport network in the secretory pathway. Represented here is the array of Golgi cisternae (blue) indexed by *j* = 1, …, *N*_*C*_ situated between the ER and PM. Glycan-binding proteins 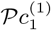 are injected from the ER to cisterna-1 at rate *q*. Superimposed is transition network of chemical reactions (column) - intercisternal transfer (rows), the latter with rates *µ*^(*j*)^. 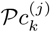 denotes the acceptor substrate in compartment *j* and the glycosyl donor *c*_0_ is chemostated in each cisterna. This results in a frequency distribution of glycans displayed at the PM (red curve), that is representative of the cell type.

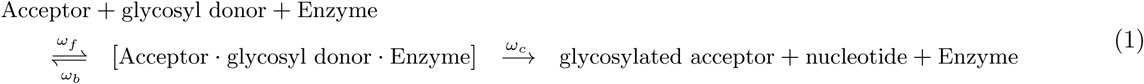

From the first cisterna, the proteins with attached sugars are delivered to the second cisterna at a given inter-cisternal transfer rate, where further chemical processing catalysed by the enzymes resident in the second cisterna occurs. This chemical processing and inter-cisternal transfer continues until the last cisterna, thereupon the fully processed glycans are displayed at the PM [2]. The network of chemical processing and inter-cisternal transfer forms the basis the physical model that we will describe next.

Any physical model of such a network of enzymatic reactions and cisternal transfer needs to be augmented by reaction and transfer rates and chemical abundances. To obtain the range of allowable values for the reaction rates and chemical abundances, we use the elaborate enzymatic reaction models, such as the KB2005 model [23–25] (with a network of 22, 871 chemical reactions and 7565 oligosaccharide structures) that predict the N-glycan distribution based on the activities and levels of processing enzymes distributed in the Golgi-cisternae of mammalian cells, and compare these predictions with N-glycan mass spectrum data. For the allowable rates of cisternal transfer, we rely on the recent study by Ungar and coworkers [26, 27], whose study shows how the overall Golgi transit time and cisternal number, can be tuned to engineer a homogeneous glycan distribution.

## IV. MODEL DEFINITION

### A. Chemical reaction and transport network in cisternae

With this background, we now define our quantitative model for chemical processing and transport in the secretory pathway. We consider an array of *N*_*C*_ Golgi cisternae, labelled by *j* = 1, …, *N*_*C*_, between the ER and the PM (Fig. 2). Glycan-binding proteins (GBPs), denoted as 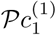, are delivered from the ER to cisterna-1 at an injection rate *q*. It is well established that the concentration of the glycosyl donor in the *j*-th cisterna is chemostated [2, 28–30], thus in our model we hold its concentration 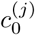 constant in time for each *j*. The acceptor 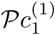 reacts with 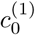 to form the glycosylated acceptor 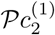, following an MM-reaction (1) catalysed by the appropriate enzyme. The acceptor 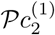 has the potential of being transformed into 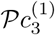, and so on, provided the requisite enzymes are present in that cisterna. This leads to the sequence of enzymatic reactions 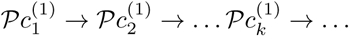, where *k* enumerates the sequence of glycosylated acceptors, using a consistent scheme (such as in [23]). The glycosylated GBPs are transported from cisterna-1 to cisterna-2 at an inter-cisternal transfer rate *µ*^(1)^, whereupon similar enzymatic reactions proceed. The processes of intra-cisternal chemical reactions and inter-cisternal transfer continue to the other cisternae and form a network as depicted in Fig. 2. Although, in this paper, we focus on a sequence of reactions that form a line-graph, the methodology we propose extends to tree-like reaction sequences, and more generally to reaction sequences that form a directed acyclic graphs [31].

Let *N*_*s*_ denote the maximum number of possible glycosylation reactions in each cisterna *j*, catalysed by enzymes labelled as 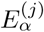, with *α* = 1, …, *N*_*E*_, where *N*_*E*_ is the total number of enzyme species in each cisterna. Since many substrates can compete for the substrate binding site on each enzyme, one expects in general that *N*_*s*_ ≫ *N*_*E*_. The configuration space of the network Fig. 2 is *N*_*s*_ *× N*_*C*_. For the N-glycosylation pathway in a typical mammalian cell, *N*_*s*_ = 2 *×* 10^4^, *N*_*E*_ = 10 − 20 and *N*_*C*_ = 4 − 8 [23–25, 27]. We account for the fact that the enzymes have specific cisternal localisation, by setting their concentrations to zero in those cisternae where they are not present.

Now the action of enzyme 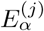 on the substrate 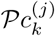 in cisterna *j* is given by

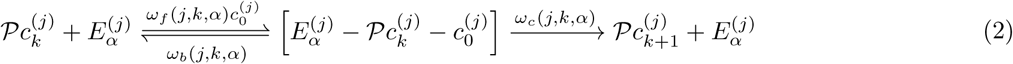

In general, the forward, backward and catalytic rates *ω*_*f*_, *ω*_*b*_ and *ω*_*c*_, respectively, depend on the cisternal label *j*, the reaction label *k*, and the enzyme label *α*, that parametrise the MM-reactions [32]. For instance, structural studies on glycosyltransferase-mediated synthesis of glycans [33], would suggest that the forward rate *ω*_*f*_ to depend on the binding energy of the enzyme 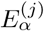 to acceptor substrate 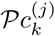 and a *physical variable that characterises cisterna*-*j*.

A potential candidate for such a cisternal variable is pH [34], whose value is maintained homeostatically in each cisterna [35]; changes in pH can affect the shape of an enzyme (substrate) or their charge properties, and in general the reaction efficiency of an enzyme has a pH optimum [32]. Another possible candidate for a cisternal variable is membrane bilayer thickness [36] - indeed both pH [37] and membrane thickness are known to have a gradient across the Golgi cisternae. We take *ω*_*f*_ (*j, k, α*) ∝ *P* ^(*j*)^(*k, α*), where *P* ^(*j*)^(*k, α*) ∈ (0, 1), is the binding probability of enzyme 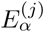 with substrate 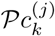, and define the binding probability *P* ^(*j*)^(*k, α*) using a biophysical model, similar in spirit to the Monod-Wyman-Changeux model of enzyme kinetics [38, 39], of enzyme-substrate induced fit.

Let 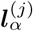 and ***l***_*k*_ denote, respectively, the optimal “shape” for enzyme 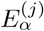 and the substrate 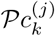. We assume that the mismatch (or distortion) energy between the substrate *k* and enzyme 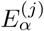 is 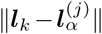, with a binding probability given by,

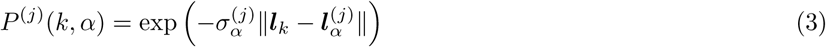

where ∥. ∥ is a distance metric defined on the space of 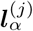 (e.g., the square of the *l*_2_-norm would be related to an elastic distortion model [40]) and the vector 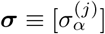 parametrises *enzyme specificity*. A large value of 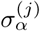 indicates a highly specific enzyme, a small value of 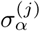 indicates a promiscuous or sloppy enzyme. It is recognised that the degree of enzyme specificity or sloppiness is an important determinant of glycan distribution [2, 41–43].

As in [23–25], our synthesis model is mean-field, in that we ignore stochasticity in glycan synthesis that may arise from low copy numbers of substrates and enzymes, multiple substrates competing for the same enzymes, and kinetics of inter-cisternal transfer. Then the usual MM-steady state condition on (2), which assumes that the concentration of the intermediate enzyme-substrate complex does not change with time, implies

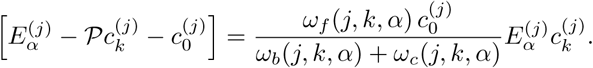

where 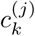 is the *concentration* of the acceptor substrate 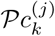 in compartment *j*.

Together with the constancy of the total enzyme concentration, 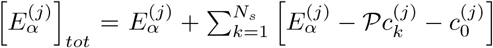, this immediately fixes the kinetics of product formation (not including inter-cisternal transport),

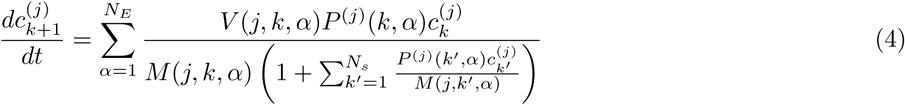

where

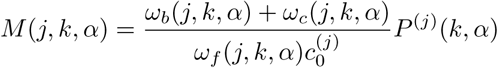

and

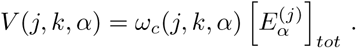

From the above, the experimentally measurable parameters *V*_*max*_ and MM-constant *K*_*M*_, for each (*j, k, α*) can be easily read out. As is the usual case, the maximum velocity *V*_*max*_ is not an intrinsic property of the enzyme, because it is dependent on the enzyme concentration 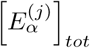 ; while 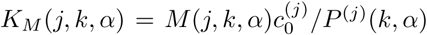 is an intrinsic parameter of the enzyme and the enzyme-substrate interaction. The enzyme catalytic efficiency, the so-called “*k*_*cat*_*/K*_*M*_” ∝ *P* ^(*j*)^(*k, α*) and is high for *perfect* enzymes [44] with minimum mismatch.

We now add to this chemical reaction kinetics, the rates of injection (*q*) and inter-cisternal transport *µ*^(*j*)^ from the cisterna *j* to *j* + 1; in Appendix A we display the complete set of equations that describe the changes in the substrate concentrations 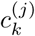 with time. These kinetic equations automatically obey the conservation law for the protein concentration (*p*). Rescaling the kinetic parameters in terms of the injection rate *q*, i.e. *V* (*j, k, α*) = *V* (*j, k, α*)*/q* and *µ*^(*j*)^ = *µ*^(*j*)^*/q*, we see that the steady state concentrations of the glycans in each cisterna satisfy the following recursion relations (see, Appendix A). In the first cisterna,

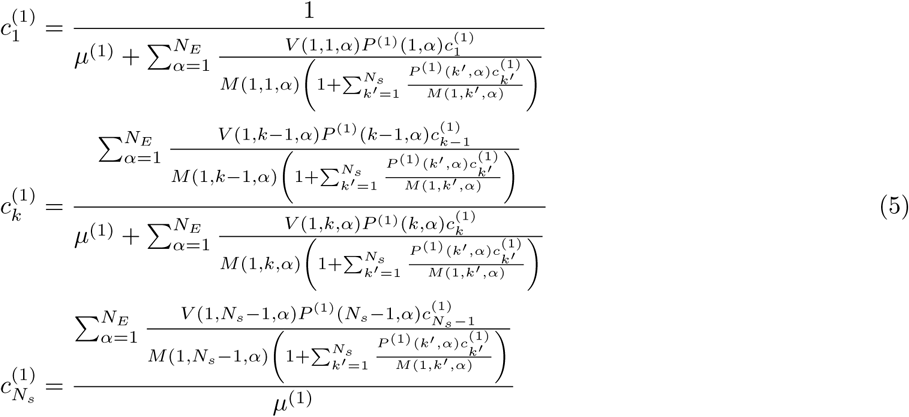

and in cisternae *j* ≥ 2,

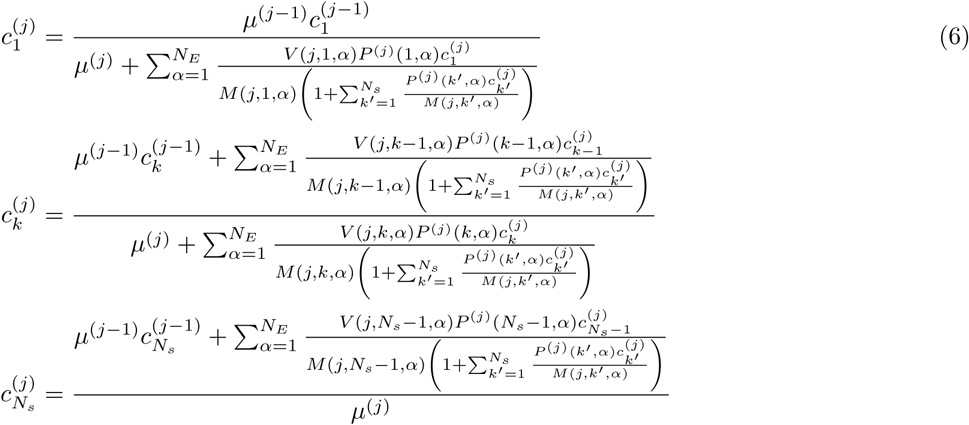

Equations (5)-(6) automatically imply that the total steady state glycan concentration in each cisterna *j* = 1, …, *N*_*c*_ is given by

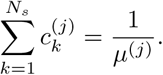

These nonlinear recursion equations (5)-(6) have to be solved numerically to obtain the steady state glycan concentrations, 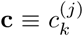, as a function of the independent vectors **M** ≡ [*M* (*j, k, α*)], **V** ≡ [*V* (*j, k, α*)], and **L** ≡ [*P* ^(*j*)^(*k, α*)], the transport rates ***µ*** ≡ [*µ*^(*j*)^] and specificity, 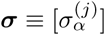.

## V. OPTIMIZATION PROBLEM

Now, with both the protocol for determining the target glycan distribution and the sequential chemical processing model in hand, we can precisely define the optimization problem referred to in the Introduction. Let **c**^*∗*^ denote the “target” concentration distribution^2^ for a particular cell type, i.e. the goal of the sequential synthesis mechanism described in Sect. IV A is to approximate **c**^*∗*^. Let 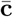 denote the steady state glycan concentration distribution displayed on the PM - (6) implies that 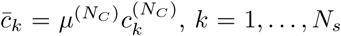. We measure the fidelity between the **c**^*∗*^ and 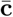 by the Kullback-Leibler metric [16, 17],

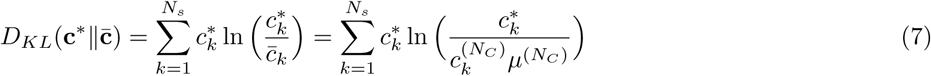

Thus, the problem of designing a sequential synthesis mechanism that approximates **c**^*∗*^ for a given enzyme specificity ***σ***, transport rate ***µ***, number of enzymes *N*_*E*_, and number of cisternae *N*_*C*_ is given by

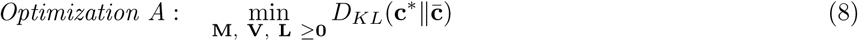

There is separation of time scales implicit in Optimization A – the chemical kinetics of the production of glycans and their display on the PM happens over cellular time scales, while the issues of tradeoffs and changes of parameters are driven over evolutionary timescales.

Optimization A, though well-defined, is a hard problem, since the steady state concentrations (6) are not *explicitly* known in terms of the parameters (**M, V, L**). In Appendix B, we formulate an alternative problem *Optimization B* in which the steady state concentrations are defined explicitly in terms of a new parameters **R** and **L**, and in Appendix C we prove that Optimization A and Optimization B are exactly equivalent. This is a crucial insight that allows us to obtain all the results that follow.

In Appendix H, we describe the variant of the Sequential Quadratic Programming (SQP) [45], that we use to numerically solve the optimization problem.

## VI. RESULTS OF OPTIMIZATION

To start with, the dimension of the optimization search space is extremely large ≈ *O*(*N*_*s*_ *× N*_*E*_ *× N*_*C*_). To make the optimization search more manageable, we ignore the *k*-dependence of the vectors (**M, V**), (or, alternatively of **R**, see Appendix B for details). The dependence on the reaction rates on the glycosyl substrate is still present in the forward reactions via the enzyme-substrate binding probability *P* ^(*j*)^(*k, α*). We further assume that shape function is a number, 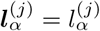 and that ***l****k* = *k*. Finally we will drop the dependence of the specificity on *α* and *j*, and take it to be a scalar *σ*. To fix our model, we will take the distortion energy that appears in (3) to be the linear form 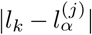. Other metrics, such as 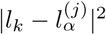, corresponding to the elastic distortion model [40], do not pose any computational difficulties, and we see that the results of our optimization remain qualitatively unchanged.

These restrictions significantly reduce the dimension of the optimization search, so much so that in certain limits, we can solve the problem analytically^3^. This helps us obtain some useful heuristics (Appendix E) on how to tune the parameters, e.g. *N*_*E*_, *N*_*C*_, *σ*, and others, in order to generate glycan distributions **c** of a given complexity. These heuristics inform our more detailed optimization using “realistic” target distributions.

The calculations in Appendix E imply, as one might expect, that the synthesis model needs to be more elaborate, i.e., needs a larger number of cisternae *N*_*C*_ or a larger number of enzymes *N*_*E*_, in order to produce a more complex glycan distribution. For a real cell type in a niche, the specific elaboration of the synthesis machinery, would depend on a variety of control costs associated with increasing *N*_*E*_ and *N*_*C*_. While an increase in the number of enzymes would involve genetic and transcriptional costs, the costs involved in increasing the number of cisternae could be rather subtle.

Notwithstanding the relative control costs of increasing *N*_*E*_ and *N*_*C*_, it is clear from the special case, that increasing the number of cisternae achieves the goal of obtaining an accurate representation of the target distribution. Let us assume that the target distribution is 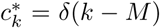 for a fixed *M* ≫ 1, i.e. 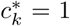 when *k* = *M*, and 0 otherwise, and that the *N*_*E*_ enzymes that catalyse the reactions are highly specific. In this limit, Optimization A reduces to a simple enumeration exercise [46]: clearly one needs *N*_*E*_ = *M*, with one enzyme species for each of the *k* = 1, …, *M* reactions, in order to generate 𝒫*c*_*M*_. For a single Golgi cisterna with a finite cisternal residence time (finite *µ*), the chemical synthesis network will generate a significant steady state concentration of lower index glycans 𝒫*c*_*k*_ with *k* < *M*, contributing to a low fidelity. To obtain high fidelity, one needs multiple Golgi cisternae with a specific enzyme partitioning (*E*_1_, *E*_2_, …, *E*_*M*_) with *E*_*j*_ enzymes in cisterna *j* = 1, …, *N*_*c*_. This argument can be generalised to the case where the target distribution is a finite sum of delta-functions. The more general case, where the enzymes are allowed to have variable specificity, needs a more detailed study, to which we turn to below.

### A. Target distribution from coarse-grained MSMS

As discussed in Sect. II, we obtain the target glycan distribution from glycan profiles for real cells obtained using Mass Spectrometry coupled with determination of molecular structure (MSMS) measurements [22]. The raw MSMS data, however, is not suitable as a target distribution. This is because it is very noisy, with chemical noise in the sample and Poisson noise associated with detecting discrete events being the most relevant [47]. This means that many of the small peaks in the raw data are not part of the signal, and one has to “smoothen” the distribution to remove the impact of noise.

We use MSMS data from *human* T-cells [22] for our analysis. As discussed in Sect. II, the Gaussian mixture models (GMM) are often used to approximate distributions with a mixed number of modes or peaks [17], or in our setting, a given fixed complexity. Here, we use a variation of the Gaussian mixture models (see Appendix G for details) to create a hierarchy of increasingly complex distributions to approximate the MSMS raw data. Thus, the 3-GMM and 20-GMM approximations represent the low and high complexity benchmarks, respectively. In Appendix G, we show that the likelihood for the glycan distribution of the *human* T-cell saturates at 20 peaks. Thus, statistically speaking, the *human* T-cell glycan distribution is accurately approximated by 20 peaks.

This hierarchy allows us to study the trade-off between the complexity of the target distribution and the complexity of the synthesis model needed to generate the distribution as follows. Let **T**^(*i*)^ denote the *i*^*th*^-GMM approximation for the *human* T-cell MSMS data. We sample this target distribution at indices *k* = 1, …, *N*_*s*_, that represent the glycan indices, and then renormalize to obtain the discrete distribution 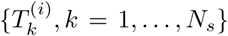. Let 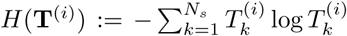 denote the entropy [16] of the *i*^*th*^-GMM approximation. *H*(**T**^(*i*)^) quantifies statistical information in the target distribution **T**^(*i*)^. We evaluate the fidelity of the distribution generated by the synthesis model to this target distribution by the ratio of the Kullback-Leibler distance to the entropy of the target distribution:

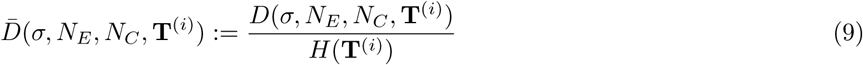

This normalization allows us evaluate the fidelity of the synthesis model to the target distribution **T**^(*i*)^ as a fraction of the total statistical information in the target distribution **T**^(*i*)^.

### B. Tradeoffs between number of enzymes, number of cisternae and enzyme specificity to achieve given complexity

We are now in a position to catalogue the main results that follow from an optimization of the parameters of the glycan synthesis machinery to a given target distribution, Figs. 3-4

**FIG. 3.**
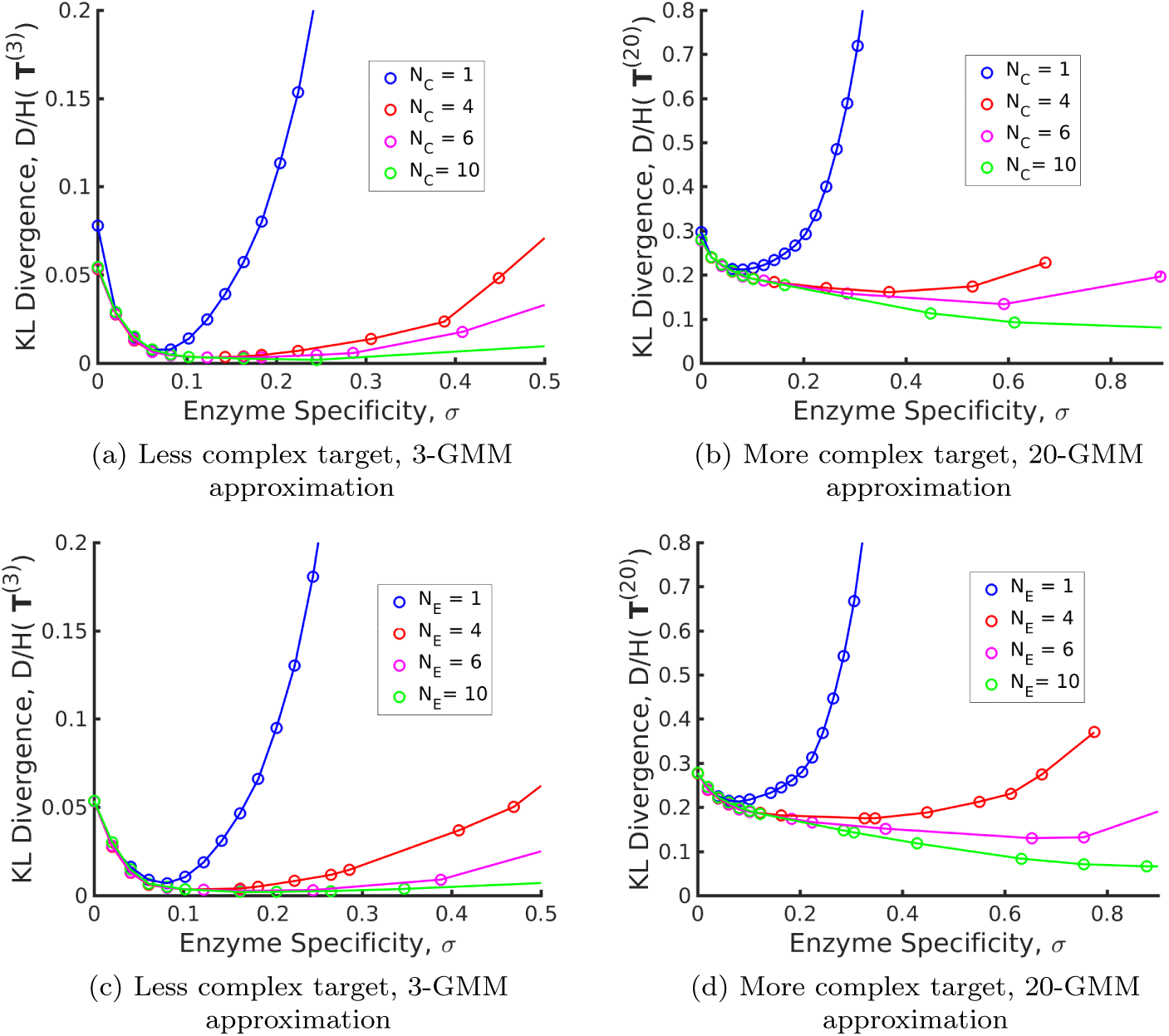
Tradeoffs amongst the glycan synthesis parameters, enzyme specificity *σ*, cisternal number *N*_*C*_ and enzyme number *N*_*E*_, to achieve a complex target distribution **c**^*∗*^). (a)-(b) Normalised Kullback-Leibler distance 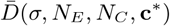 as function of *σ* and *N*_*C*_ (for fixed *N*_*E*_ = 3), (c)-(d) 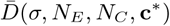 as function of *σ* and *N*_*E*_ (for fixed *N*_*C*_ = 3), with the target distribution **c**^*∗*^ set to the 3-GMM (less complex) and 20-GMM (more complex) approximations for the *human* T-cell MSMS data. 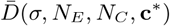 is a convex function of *σ* for each (*N*_*E*_, *N*_*C*_, **c**^*∗*^), decreasing in *N*_*C*_, *N*_*E*_ for each (*σ*, **c**^*∗*^), increasing in the complexity of **c**^*∗*^ for fixed (*σ, N*_*E*_, *N*_*C*_). The specificity 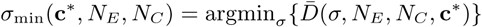 that minimises the error for given (*N*_*E*_, *N*_*C*_, **c**^*∗*^) is an increasing function of *N*_*C*_, *N*_*E*_ and the complexity of the target distribution **c**^*∗*^. Furthermore, the curvature of 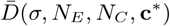 at *σ*_min_(*N*_*E*_, *N*_*C*_, **c**^*∗*^), related to *sensitivity*, is a decreasing function of *N*_*C*_, *N*_*E*_.

**FIG. 4.**
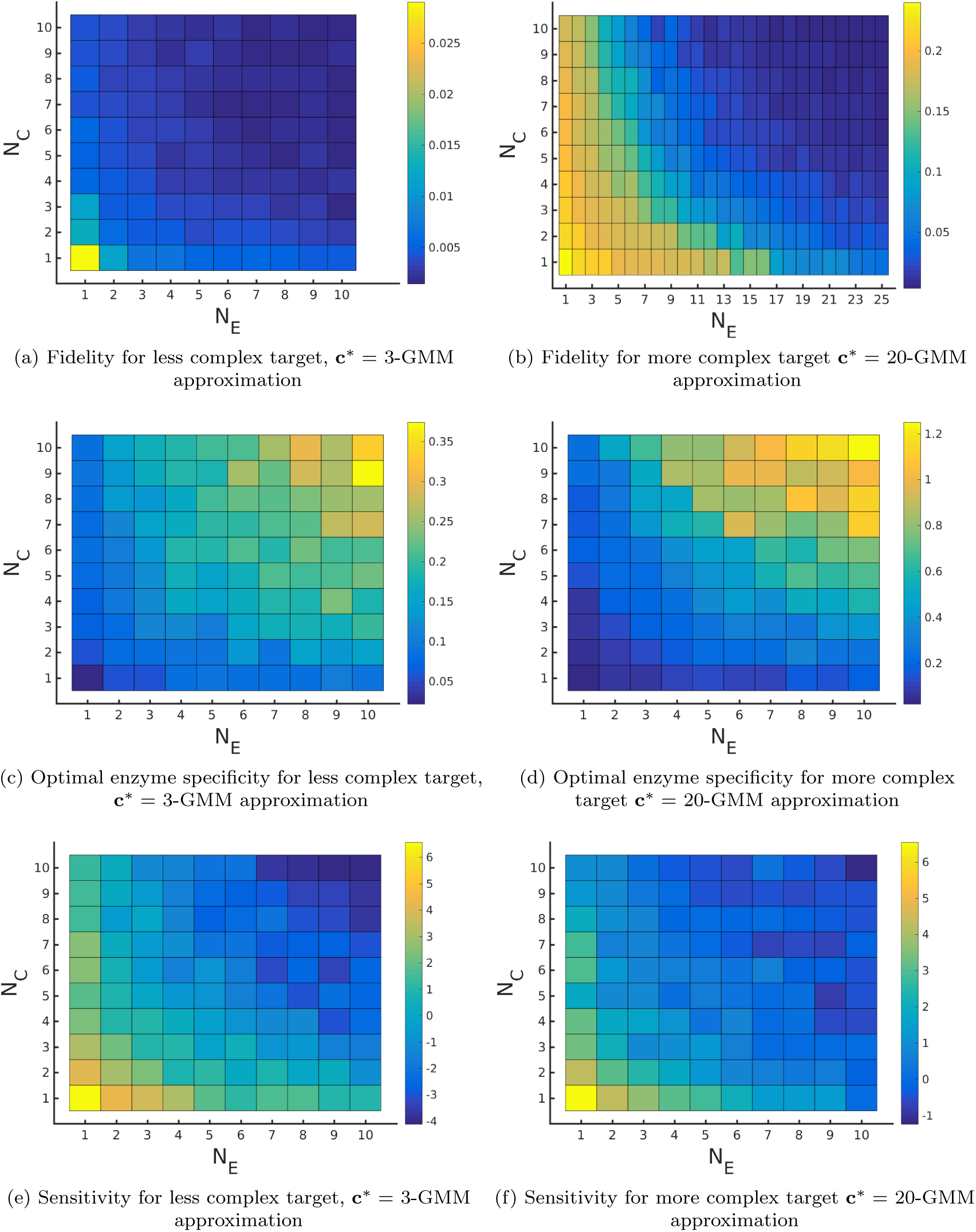
Fidelity of glycan distribution and optimal enzyme properties to achieve a complex target distribution. The target **c**^*∗*^ is taken from 3-GMM (less complex) and 20-GMM (more complex) approximations of the *human* T-cell MSMS data. (a)-(b) Minimum normalised KL divergence 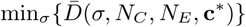 as a function of (*N*_*E*_, *N*_*C*_). More complex distributions require either a larger value *N*_*E*_ or *N*_*C*_. The marginal impact of increasing *N*_*E*_ and *N*_*C*_ on 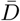 is approximately equal. (c)-(d) Optimum enzyme specificity *σ*_min_ obtained from 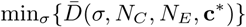 as a function of (*N*_*E*_, *N*_*C*_). *σ*_min_ increases with increasing *N*_*E*_ or *N*_*C*_. To synthesize the more complex 20 GMM approximation with high fidelity requires enzymes with higher specificity *σ*_min_ compared to those needed to synthesize the broader, less complex 3-GMM approximation. (e) -(f) Sensitivity 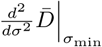 of the normalised Kullback-Leibler distance 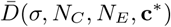 as a function of (*N*_*E*_, *N*_*C*_). Increasing *N*_*E*_ or *N*_*C*_ decreases this sensitivity implying the specificity *σ* does not need to be tuned very carefully if *N*_*E*_, *N*_*C*_ are high.

1. The normalized KL-distance 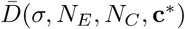 is a convex function of *σ* for fixed values for other parameters (Fig. 3), i.e. it first decreases with *σ* and then increases beyond a critical value of *σ*_min_. 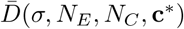 is decreasing in *N*_*C*_ and *N*_*E*_ for fixed values of the other parameters, and increasing in the complexity of **c**^*∗*^ for fixed (*σ, N*_*C*_). The marginal contribution of *N*_*C*_ and *N*_*E*_ in reducing the normalised distance 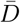 is approximately equal (Figs. 4a, 4b). The lower complexity distributions can be synthesized with high fidelity with small (*N*_*E*_, *N*_*C*_), whereas higher complexity distributions require significantly larger (*N*_*E*_, *N*_*C*_), Figs. 4a, 4b. For a typical mammalian cell, the number of enzymes in the N-glycosylation pathway are in the range *N*_*E*_ = 10−20 [23–25, 27], Fig. 4b would then suggest that the optimal cisternal number would range from *N*_*C*_ = 3 − 8 [18].
2. The optimal enzyme specificity 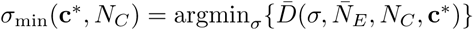, that minimises the error as function of (*N*_*C*_, **c**^*∗*^) with *N*_*E*_ fixed at 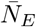, is an increasing function of *N*_*C*_ and the complexity of the target distribution **c**^*∗*^ (Figs. 3a, 3b, 4c, 4d). This is consistent with the results in Appendix E where we established that the width of the synthesized distribution is inversely dependent on the specificity *σ*: since a GMM approximation with fewer peaks has wider peaks, *σ*_min_ is low, and vice versa. Similar results hold when *N*_*C*_ is fixed at 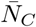, and *N*_*E*_ is varied (Figs. 3c, 3d, 4c, 4d).
3. Let *σ*_min_(*N*_*C*_, *N*_*E*_, **c**^*∗*^) denote the value of *σ* that minimizes 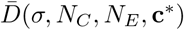. Then the second-derivative 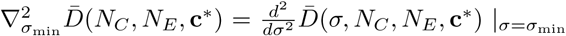 denotes the curvature at *σ* min, and is measure of the sensitivity of 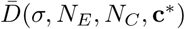 to *σ* for values close to *σ*_min_(*N*_*E*_, *N*_*C*_, **c**^*∗*^). 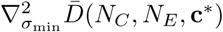 is a decreasing function of *N*_*C*_ (resp. *N*_*E*_) for fixed values of (*N*_*E*_, **c**^*∗*^) (resp. (*N*_*C*_, **c**^*∗*^)), see Figs. 3, 4e, 4f. Thus, for any target distribution **c**^*∗*^ there is a minimal value of (*N*_*E*_, *N*_*C*_) such that the target can be synthesized with high fidelity provided the sensitivity *σ* is tightly controlled at *σ*_min_(*N*_*C*_, *N*_*E*_, **c**^*∗*^), and there is larger value of (*N*_*E*_, *N*_*C*_) such that the target can be synthesized even if the control on *σ* is less tight.

Ungar et al. [26] optimize incoming glycan ratio, transport rate and effective reaction rates in order to synthesize a narrow target distribution centred around a desired glycan. The ability to produce specific glycans without much heterogeneity is an important goal in pharma industry. They define heterogeneity as the total number of glycans synthesized, and show that increasing the number of compartments *N*_*C*_ decreases heterogeneity, and increases the concentration of the specific glycan. They also show that changing transport rate does not affect the heterogeneity. Our results are entirely consistent with theirs - we have shown that 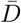 decreases as we increase *N*_*C*_. Thus, if the target distribution has a single sharp peak, increasing *N*_*C*_ will reduce the heterogeneity in the distribution.

### C. Optimal partitioning of enzymes in cisternae

Having studied the optimum *N*_*E*_, *N*_*C*_, *σ* to attain a given target distribution with high fidelity, we ask what is the optimal partitioning of the *N*_*E*_ enzymes in these *N*_*C*_ cisternae? Answering this within our chemical reaction model (Sect. IV A) requires some care, since it incorporates the following enzymatic features: (a) enzymes with a finite specificity *σ* can catalyse several reactions, although with an efficiency that varies with both the substrate index *k* and cisternal index *j*, and (b) every enzyme appears in each cisternae; however their reaction efficiencies depend on the enzyme levels, the enzymatic reaction rates and the enzyme matching function **L**, all of which depend on the cisternal index *j*.

Thus, rather than determining the cisternal partitioning of enzymes, we instead identify chemical reactions that occur with high propensity in each cisternae. For this we define an effective reaction rate 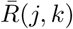 for *𝒫 c*_*k*_ → *𝒫 c*_*k*+1_ in the *j-*th cisterna as

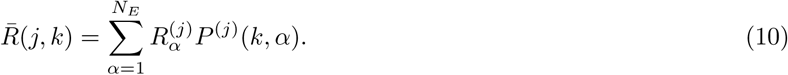

According to our model presented in Sect. IV A, the list of reactions with high effective reaction rates in each cisterna, corresponds to a cisternal partitioning of the perfect enzymes. In a future study, we will consider a Boolean version of a more complex chemical model, to address more clearly, the optimal enzyme partitioning amongst cisternae.

Figure 5 (a) (i) shows the heat map of the effective reaction rates in each cisterna for the optimal *N*_*E*_, *N*_*C*_, *σ* that minimises the normalised KL-distance to the 20 GMM target distribution **T**^(20)^ (see Fig. 5 (a) (ii)). The optimized glycan profile displayed in Fig. 5 (a) (iii) is very close to the target. An interesting observation from Fig. 5a(i) is that the same reaction can occur in multiple cisternae.

**FIG. 5.**
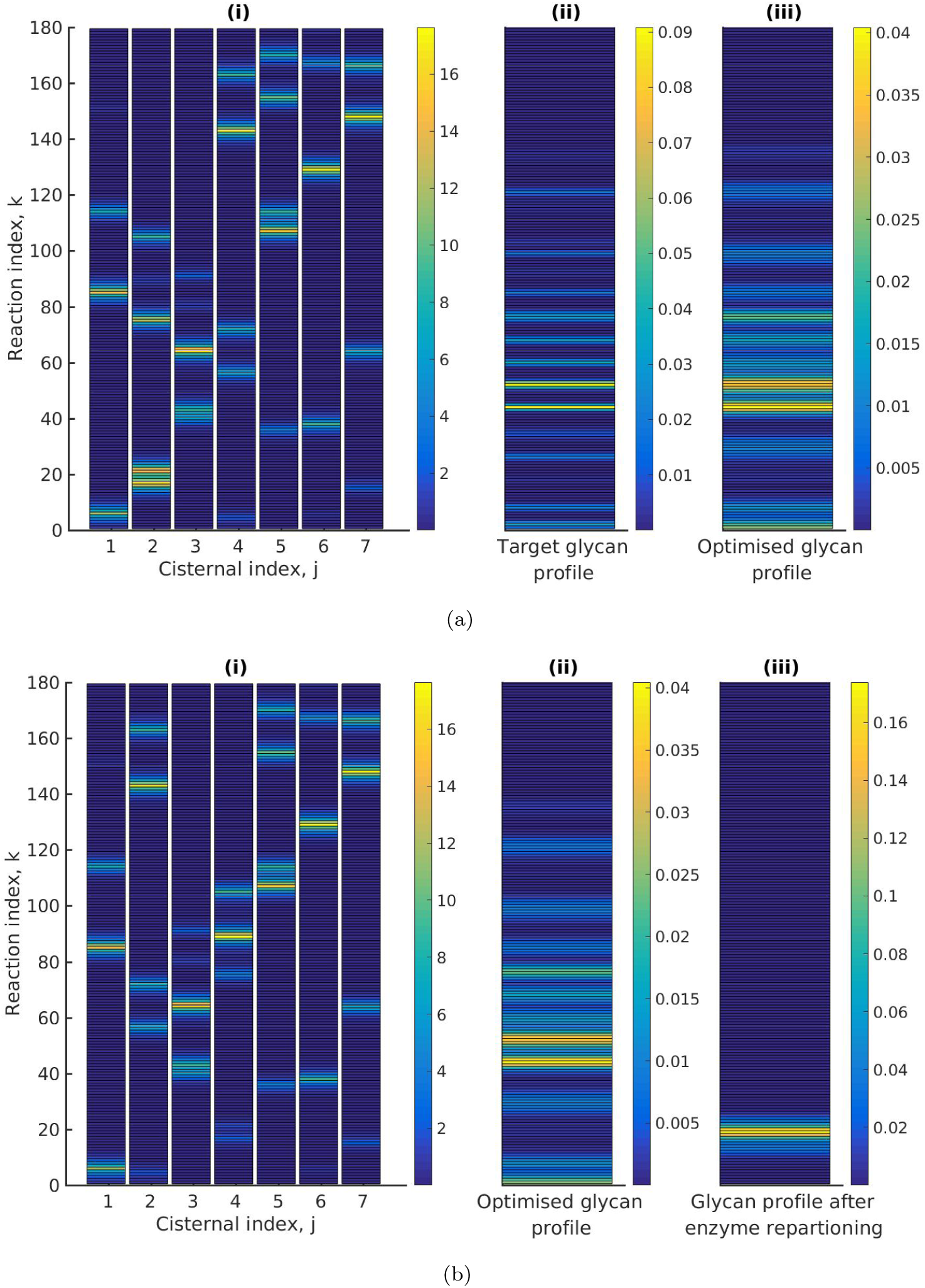
Optimal enzyme partitioning in cisternae. (a) Heat map of the effective reaction rates in each cisterna (representing the optimal enzyme partitioning) and the steady state concentration in the last compartment 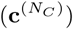 for the 20-GMM target distribution. Here *N*_*E*_ = 5, *N*_*C*_ = 7, normalised 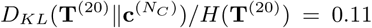. (b) Effective Reaction rates after swapping the optimal enzymes of the fourth and second cisternae. The displayed glycan profile is considerably altered from the original profile.

Keeping everything else fixed at the optimal value, we ask whether simply repartitioning the optimal enzymes amongst the cisternae, alters the displayed glycan distribution. In Fig. 5 (b) (i), we have exchanged the enzymes of the fourth and second cisterna. The glycan profile after enzyme partitioning (see Fig. 5 (b) ((iii)) is now completely altered (compare Fig. 5 (b) (ii) with Fig. 5 (b) (iii)). Thus one may achieve a different glycan distribution by repartitioning enzymes amongst the same number of cisternae [46].

## VII. STRATEGIES TO ACHIEVE HIGH GLYCAN DIVERSITY

So far we have studied how the complexity of the target glycan distribution places constraints on the evolution of Golgi cisternal number and enzyme specificity. We now take up another issue, namely, how the physical properties of the Golgi cisternae, namely cisternal number and inter-cisternal transport rate, may drive diversification of glycans [48, 49]. There is substantial correlative evidence to support the idea that cell types that carry out extensive glycan processing employ larger numbers of Golgi cisternae. For example, the salivary Brunners gland cells secrete mucous that contains heavily O-glycosylated mucin as its major component [50]. The Golgi complex in these specialized cells contain 9 − 11 cisternae per stack. Additionally, several organisms such as plants and algae secrete a rather diverse repertoire of large, complex glycosylated proteins, for a variety of functions [51–60]. These organisms possess enlarged Golgi complexes with multiple cisternae per stack [61–65].

In this section, we study how changing the physical parameters in our chemical synthesis model can lead to changes in the diversity of glycan distributions.

We define *diversity* as the total number of glycan species produced above a specified threshold abundance *c*_*th*_. This last condition is necessary because very small peaks will not be distinguishable in the presence of noise. In computing the diversity from our chemical synthesis model, we have chosen the threshold to be *c*_*th*_ = 1*/N*_*s*_, where *N*_*s*_ is the total number of glycan species. We have checked that the qualitative results do not depend on this choice, Fig. A6.

Using the sigmoid function (1 + *e*^−*x/τ*^)^−1^ as a continuous approximation to the Heaviside function Θ(*x*), we define the following optimization to achieve the maximal diversity for a given set of parameter values, *N*_*E*_, *N*_*C*_, *σ*,

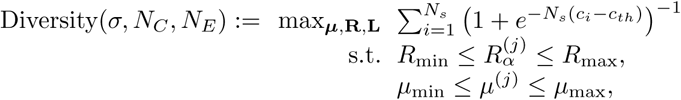

where, as before, (*µ*_max_, *µ*_min_) = (1, 0.01)/min, and (*R*_max_, *R*_min_) = (20, 0.018)/min, and *c*_*th*_ = 1*/N*_*s*_ is the threshold. See Appendix F for details on the parameter estimation.

The results displayed in Fig. 6(a), show that for a fixed specificity *σ*, the diversity at first increases with the number of cisternae *N*_*C*_, and then saturates at a value that depends on *σ*. For very high specificity enzymes, one can achieve very high diversity by appropriately increasing *N*_*C*_. This establishes the link between glycan diversity and cisternal number. However, this link is correlative at best, since there are many ways to achieve high glycan diversity - notably by increasing the number of enzymes.

**FIG. 6.**
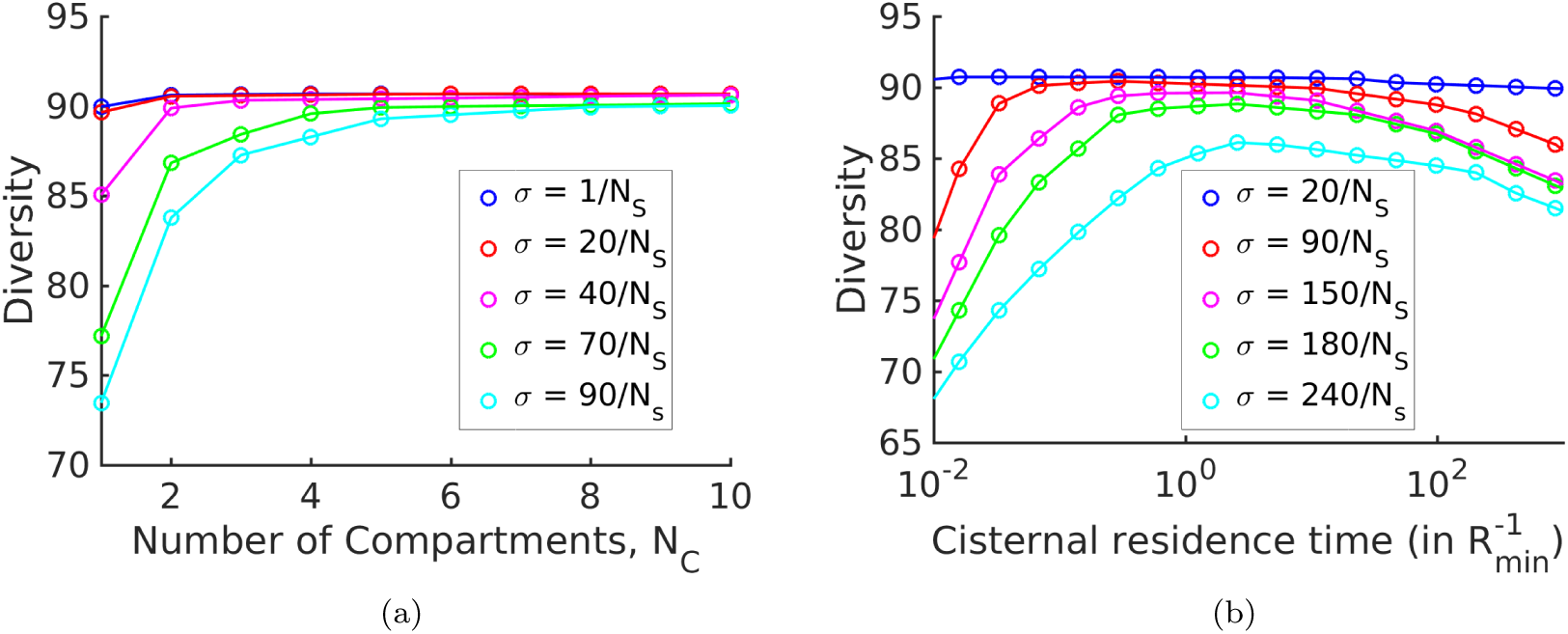
Strategies for achieving high glycan diversity. Diversity versus *N*_*C*_ and transport rate *µ* at various values of specificity *σ* for fixed *N*_*E*_ = 3. (a) Diversity vs. *N*_*C*_ at optimal transport rate *µ*. Diversity initially increases with *N*_*C*_, but eventually levels off. The levelling off starts at a higher *N*_*C*_ when *σ* is increased. These curves are bounded by the *σ* = 0 curve. (b) Diversity vs. cisternal residence time (*µ*^−1^) in units of the reaction time 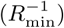 at various value of *σ*, for fixed *N*_*C*_ = 4 and *N*_*E*_ = 10. This has implications for glycoengineering (see text) where the task is to produce a particular glycan profile with low heterogeneity [26, 46].

On the other hand, one of the goals of glycoengineering is to produce a particular glycan profile with low heterogeneity [26, 46]. For low specificity enzymes, the diversity remains unchanged upon increasing the cisternal residence time. For enzymes with high specificity, the diversity typically shows a non-monotonic variation with the cisternal residence time. At small cisternal residence time, the diversity decreases from the peak because of early exit of incomplete oligomers. At large cisternal residence time the diversity again decreases as more reactions are taken to completion. Note that the peak is generally very flat, this is consistent with the results of [26]. To get a sharper peak, as advocated for instance by [46], one might need to increase the number of high specificity enzymes *N*_*E*_ further.

## VIII. DISCUSSION

The precision of the stereochemistry and enzymatic kinetics of these N-glycosylation reactions [2], has inspired a number of mathematical models [23–25] that predict the N-glycan distribution based on the activities and levels of processing enzymes distributed in the Golgi-cisternae of mammalian cells, and compare these predictions with N-glycan mass spectrum data. Models such as the KB2005 model [23–25] are extremely elaborate (with a network of 22, 871 chemical reactions and 7565 oligosaccharide structures) and require many chemical input parameters. These models have an important practical role to play, that of being able to predict the impact of the various *chemical parameters* on the glycan distribution, and to evaluate appropriate metabolic strategies to recover the original glycoprofile. Additionally, a recent study by Ungar and coworkers [26, 27] shows how *physical parameters*, such as overall Golgi transit time and cisternal number, can be tuned to engineer a homogeneous glycan distribution. Overall, such models may help predict glycosylation patterns and direct glycoengineering projects to optimize glycoform distributions.

In this paper, we have been interested in the role of glycans as a marker or molecular code of cell identity [4, 7, 11]. In particular, we have studied one aspect of molecular coding, namely the *fidelity* of this glycan code generated by enzymatic and transport processes located in the secretory apparatus of the cell. This involves a method of analysis that draws on many different fields, and so it might be useful to provide a short summary of the assumptions, methods and results of the paper:

1. The distribution of glycans at the cell surface is a marker of *cell-type identity* [2, 4, 7, 11]. This glycan distribution can be very complex; it is believed that there is an evolutionary drive for having glycan distributions of high *complexity* arising from the following considerations,
  a. Reliable cell type identification amongst a large set of different cell types in a complex organism, the preservation and diversification of “self-recognition” [5].
  b. Consequence of pathogen-mediated selection pressures [2, 4, 6].
  c. Consequence of *herd immunity* within a heterogenous population of cells of a community [15] or within a single organism [5].
2. The glycans at the cell surface are the end product of a sequential chemical processing via a set of enzymes resident in the Golgi cisternae, and transport across cisternae [4, 10, 11]. Using a fairly general and tractable model for chemical synthesis and transport, we compute the *synthesized* glycan distribution at the cell surface. Parameters of our synthesis model include the number of enzymes *N*_*E*_, specificity of enzymes *σ*, number of cisternae *N*_*C*_ and transport rates *µ*.
3. We measure the reliability or fidelity of cell identity [10, 11, 66] in terms of the error between synthesized glycan distribution on the cell surface from the its internal “target” distribution using the Kullback-Leibler distance *D*_*KL*_ [16, 17]. In our numerical study, we obtain the *target distribution* for the given cell type by suitable coarse-graining of the MSMS data for the *human* T-cells [22]. We solve a constrained optimization problem for minimising *D*_*KL*_, and study the tradeoffs between *N*_*E*_, *N*_*C*_ and *σ*.
4. The results of the optimization to achieve a given target complexity are summarised in Figs. 3-4. Here, we highlight some its direct consequences:
  a. Keeping the number of enzymes fixed, a more elaborate transport mechanism (via control of *N*_*C*_ and *µ*) is essential for synthesising high complexity target distributions (Figs. 4a, 4b). Fewer cisternae cannot be compensated for by optimising the enzymatic synthesis (via control of parameters **R, L** and *σ*).
  b. Thus, our study suggests that fidelity of the glycan code generation provides a functional control of Golgi cisternal number. It also provides a quantitative argument for the evolutionary requirement of multiple-compartments, by demonstrating that the fidelity and sensitivity of the glycan code arising from a chemical synthesis that involves multiple cisternae is higher than one that involves a single cisterna (keeping everything else fixed) (Figs. 4a, 4b, 4e, 4f). This feature that with multiple cisternae and precise enzyme partitioning, one may generically achieve a highly accurate representation of the target distribution, has been highlighted in an algorithmic model of glycan synthesis [46].
  c. Our study shows that for a fixed *N*_*C*_ and *N*_*E*_, there is an optimal enzyme specificity that achieves the lowest distance from a given target distribution. As we see in Fig. 4d, this optimal enzyme specificity can be very high for highly complex target distributions.
  d. Organisms such as plants and algae, have a diverse repertoire of glycans that are utilised in a variety of functions [51–60]. Our study shows that it is optimal to use low specificity enzymes to synthesize target distributions with high diversity (Fig. 6). However, this compromises on the complexity of the glycan distribution, revealing a tension between complexity and diversity. One way of relieving this tension is to have larger *N*_*E*_ and *N*_*C*_.
  e. Consider a situation where the environment, and hence the target glycan distribution, fluctuates rapidly. When synthesis parameters cannot change rapidly enough to track the environment, high specificity enzymes can lead to a *lowering* of the cell’s fitness [67, 68]. Having slightly sloppy enzymes may give the best selective advantage in a time varying environment. This compromise, between robustness in a changing environment and the demand for complexity, is achieved by having sloppy enzymes, that allows the system to be more *evolvable* [67, 68]. However, sloppy enzymes are subject to errors from synthesising the wrong reaction products. In this case, error correcting mechanisms must be in place to ensure fidelity of the glycan code. We leave the role of intra-cellular transport in providing non-equilibrium proof-reading mechanisms to reduce such coding errors, and its optimal adaptive strategies and plasticity in a time varying environment, as a task for the future.

Admittedly the chemical network that we have considered here is much simpler than the chemical network associated with all possible protein modifications in the secretory pathway. For instance, typical N-glycosylation pathways would involve the glycosylation of a variety of GBPs. Further, apart from N-glycosylation, there are other glycoprotein, proteoglycan and glycolipid synthesis pathways [1, 2, 11]. We believe our analysis is generalisable and that the qualitative results we have arrived at would still hold.

To conclude, our work establishes the link between the cisternal machinery (chemical and transport) and optimal coding. We find that the pressure to achieve the target glycan code for a given cell type, places strong constraints on the cisternal number and enzyme specificity [18]. An important implication is that a description of the nonequilibrium self-assembly of a fixed number of Golgi cisternae must combine the dynamics of chemical processing and membrane dynamics involving fission, fusion and transport [18–20]. We believe this is a promising direction for future research.

## IX. ACKNOWLEDGMENTS

We thank M. Thattai, A. Jaiman, S. Ramaswamy, A. Varki for discussions, and S. Krishna and R. Bhat for very useful suggestions on the manuscript. We thank our group members at the Simons Centre for many incisive inputs. We are very grateful to P. Babu for consultations on the MSMS data and literature. We acknowledge the computational facilities at NCBS. MR acknowledges a JC Bose Fellowship from DST (Government of India), and thanks Institut Curie for hosting a visit under the Labex program. This work has received support under the program Investissements dAvenir launched by the French Government and implemented by ANR with the references ANR-10-LABX-0038 and ANR-10-IDEX-0001-02 PSL. QV thanks the Simons Centre (NCBS) for hosting his visit.

## Appendix

## Appendix A: Kinetics of sequential chemical reactions and transport

On including the rates of injection (*q*) and inter-cisternal transport *µ*^(*j*)^ from the cisterna *j* to *j* + 1, into the chemical reaction kinetics, the substrate concentrations 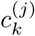 change with time as,

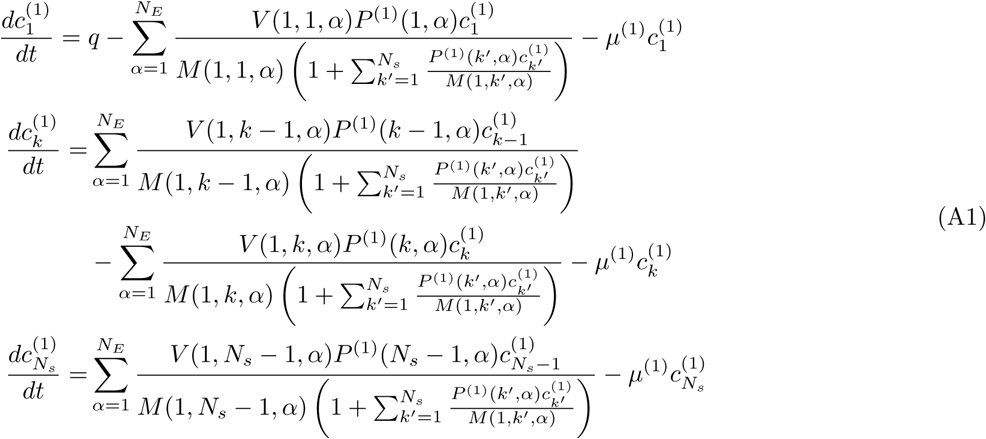

for cisterna-1, and

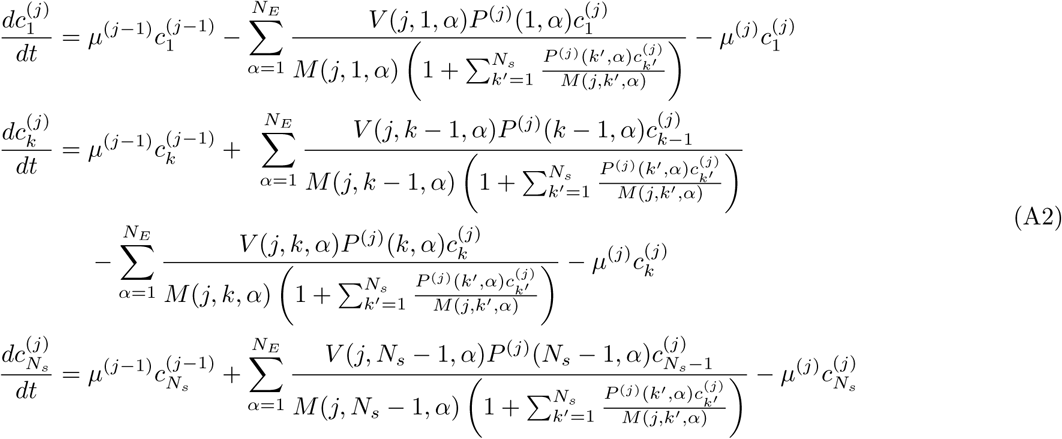

for cisternae *j* = 2, 3, …, *N*_*C*_. These set of dynamical equations (A1)-(A2), with initial conditions, can be solved to obtain the concentration 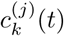 for *t* ≥ 0.

Equations (A1)-(A2) automatically obey the conservation law for the protein concentration (*p*), i.e., denoting the protein concentration in the *j*-th cisterna as 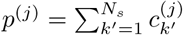, we automatically obtain,

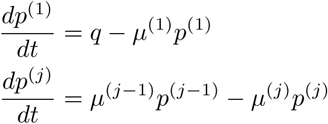

for *j* = 2, 3, … *N*_*C*_.

At steady state, the left hand side of the above equations is set to zero, which after rescaling, gives the nonlinear recursion relations displayed in (5) and (6) of the main text.

## Appendix B: A computationally simpler optimization equivalent to Optimization A

Define a new set of parameters,

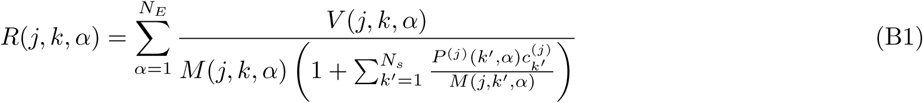

where **c** denotes the steady state glycan concentration, corresponding to a specific (**M, V, L**). Define **v** by the following set of linear equations:

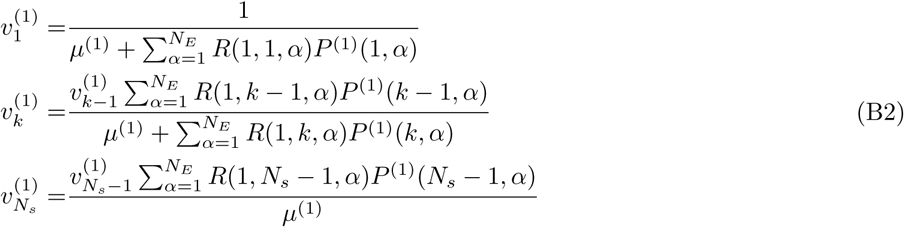

for *j* = 1, and

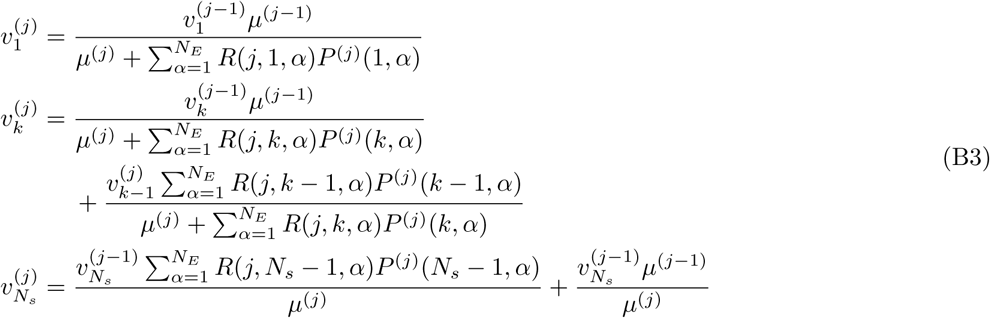

for *j* = 2, …, *N*_*C*_. Then, by the definition of **R** in (B1), it trivially follows that the steady state concentration **c** corresponding to (**M, V, L**) is a solution for (B2)-(B3).

In Appendix C we show that for **v** obtained from (B2)-(B3) for any parameter (**R, L**), there exists parameter (**M, V, L**) such that (5)-(6) are automatically satisfied when we set **c** = **v**, i.e. **v** is the steady state concentration for (**M, V, L**). Thus, the set of all concentration profiles defined by (B2)-(B3) as a function of all possible values of the parameters (**R, L**) is identical to the set defined by (5)-(6) as function of (**M, V, L**). This is a crucial insight, since it allows us to search the entire parameter space using (B2)-(B3), where the concentration is known explicitly in terms of (**R, L**). See Figure A1 for a flow chart of the two optimization schemes.

To pose this new optimization problem, it is convenient to define 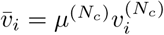. Then, it follows that *Optimization B*:

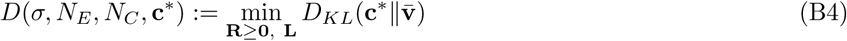

**FIG. A1.**
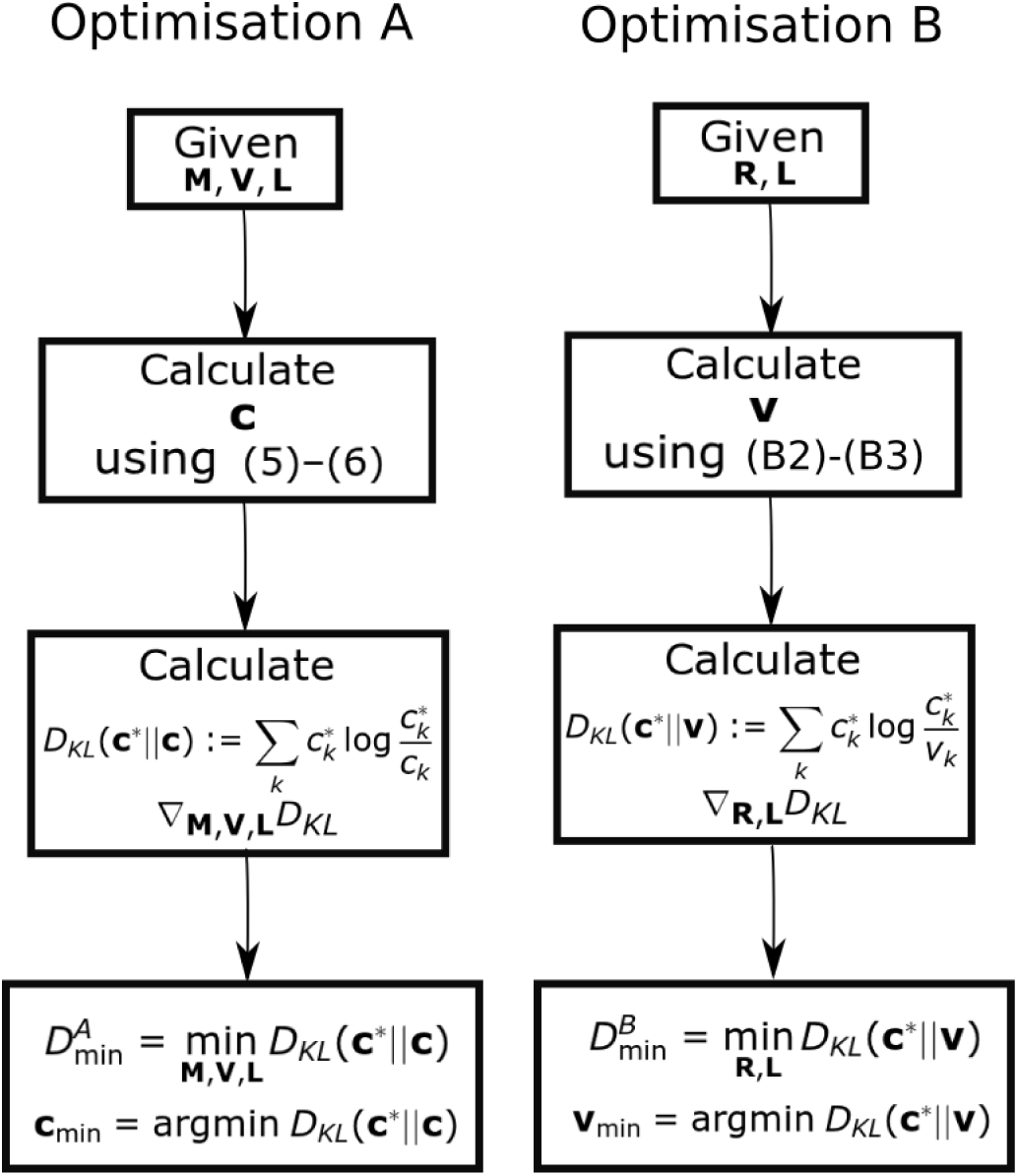
Flow chart showing the optimization schemes for Optimization A and B. We prove that 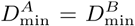 by showing the set of all **c** is equal to the set of all **v**. We additionally establish that the optimum **v**_min_ = **c**_min_.

is equivalent to (8). Since **v** is explicitly known as a function of (**R, L**), optimization B (B4) is a more tractable optimization problem than (8). Note that in this setting, the function *D*(*σ, N*_*E*_, *N*_*C*_, **c**^*∗*^) (B4) is independent of the rates ***µ***.

While this optimization is easy to implement, we note that the parameters (e.g., reaction rates, specificity) are not constrained to take only physically relevant values; a legitimate concern is that the absence of such physico-chemical constraints might drive this optimization to physically unrealistic solutions.

There are two possible ways to impose these parameter constraints. One is to impose constraints on the “microscopic” chemical parameters, such as the rate of individual reactions *R*(*j, k, α*) and the inter-cisternal transport rate *µ*^(*j*)^. These take into consideration constraints arising from molecular enzymatic processes. The other is to impose constraints on “global” physical parameters, such as the total transport time across the Golgi cisternae and the average enzymatic reaction time. Here, we impose constraints on the microscopic reaction and transport parameters.

*Optimization C* :

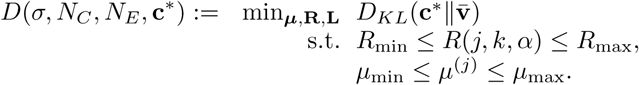

The upper and lower bounds on the rates **R** and ***µ*** are estimated in Appendix F : *µ*_max_ = 1/min (resp. *µ*_min_ = .01/min) and *R*_max_ = 20/min (resp. *R*_min_ = .018/min).

## Appendix C: Equivalence of Optimizations *A* and *B*

Let

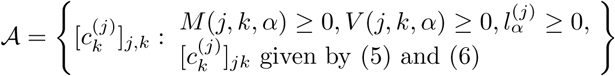

denote the set of concentrations achievable in Optimization *A*. Similarly, let

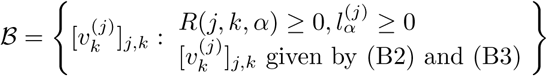

denote the set of concentrations achievable in the Optimization *B*.

Our task is to show that 𝒜 = ℬ. Suppose 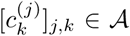. Let [*M* (*j, k, α*)], [*V* (*j, k, α*)] and 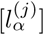 be the corresponding parameters. Define

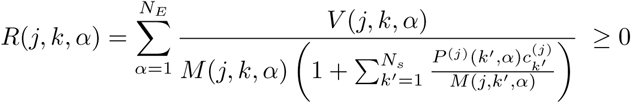

Then 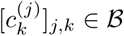.

Suppose 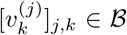. Let 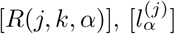 denote the corresponding parameters. Since 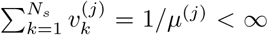, it follows that 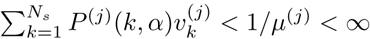. Thus, there exists parameters [*M* (*j, k, α*)], [*V* (*j, k, α*)] and 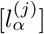 such that

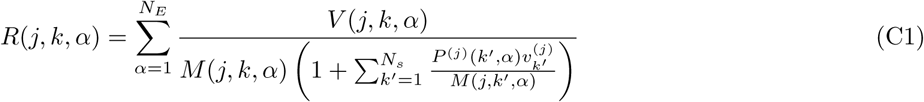

Therefore, 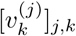 satisfy (5) and (6), i.e. 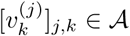.

Moreover, suppose **v** satisfies (B2)-(B3) for a given set of parameters (**R, L**). Then there exist (**M, V, L**) such that **v** satisfies (5)-(6), i.e. **v** is the steady state concentration for (**M, V, L**).

## Appendix D: Analytical solution when *N*_*s*_ ≫ 1

It is possible to obtain analytical expressions for the steady state glycan distribution, in the limit *N*_*s*_ ≫ 1, when the glycan index *k* can be approximated by a continuous variable. In this case, (5)-(6) can be cast as differential equations,

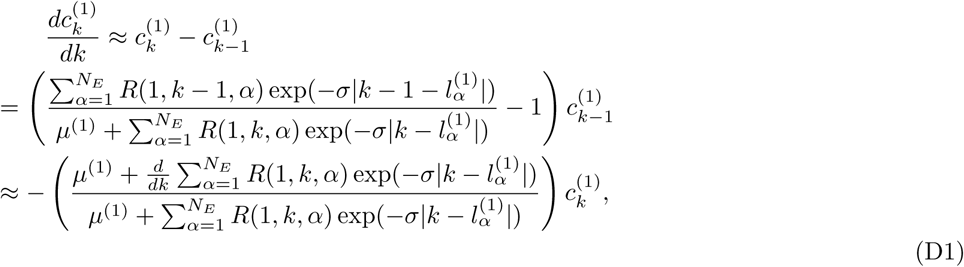

and

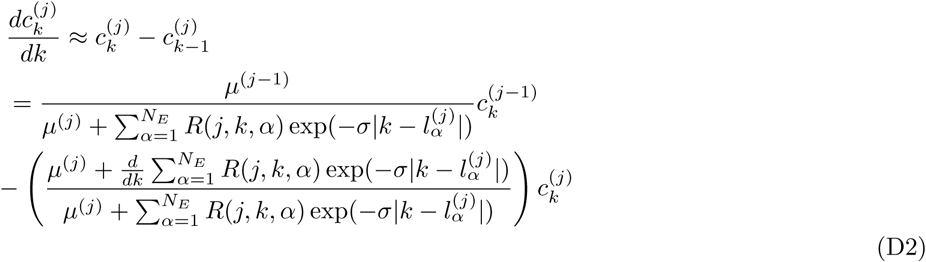

for *j* = 2, …, *N*_*C*_. In (D1) and (D2),

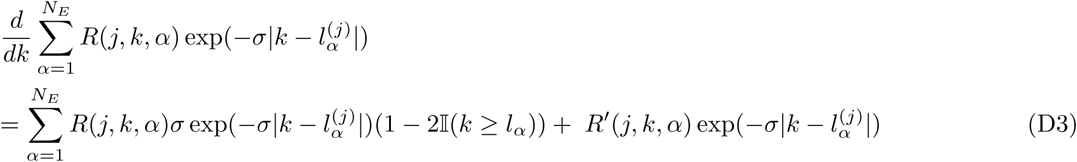

where the indicator function 𝕀(·) is equal to 1 if the argument is true, and zero otherwise and *R*′(*j, k, α*) is the derivative of *R*(*j, k, α*) with respect to *k*.

Define a vector function 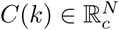 of the continuous variable *k* by 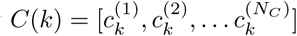. Then (D1) and (D2) can be written as:

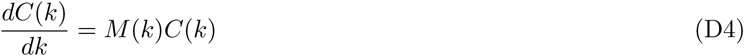

where the matrix *M* (*k*) is given by

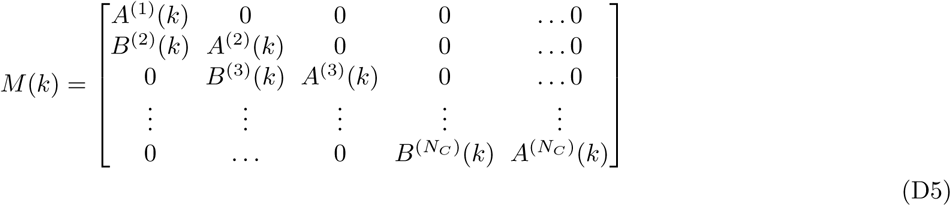

with

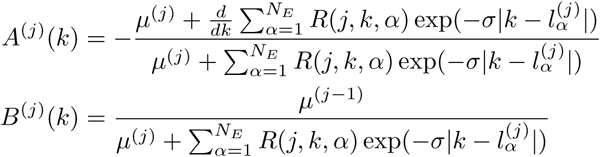

The functions *A*^(*j*)^(*k*) and *B*^(*j*)^(*k*) involve absolute value and indicator functions; therefore, the differential equation has to be solved in a piecewise manner assuming continuity of solution *C*(*k*).

The general solution of (D4)

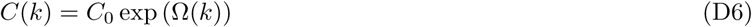

is written in terms of the Magnus Function 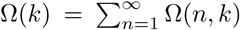, obtained from the Baker-Campbell-Hausdorff formula [69],

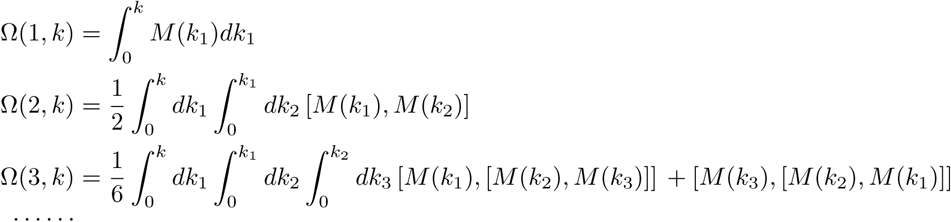

where [*M* (*k*_1_), *M* (*k*_2_)] := *M* (*k*_1_)*M* (*k*_2_) − *M* (*k*_2_)*M* (*k*_1_) is the commutator, and the higher order terms in … contain higher order nested commutators.

Here, we establish conditions under which the series 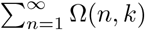 that defines solution *C*(*k*) to the differential equation (D4) converges. We also solve (D4) for some special cases.

The commutator

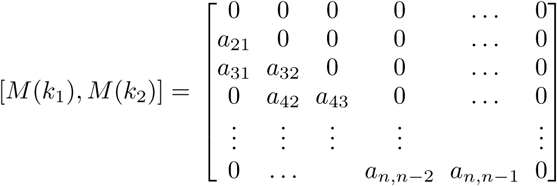

where

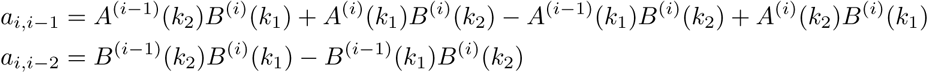

The general form of Ω(*n, k*) is given by [69]

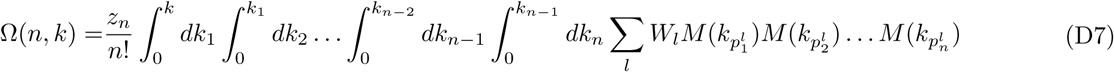

where 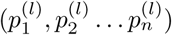 is a permutation of (1, 2, 3, … *n*), *W*_*l*_ ∈ {−1, 1}, and *z*_*n*_ ∈ 1, …*n*.

Let *Ā* = max_*k,l,m*_ |*M*_*l,m*_(*k*) |. Define

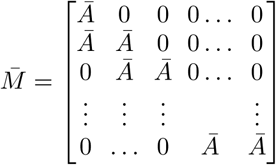

We can bound all the matrix elements of Ω(*n, k*) in the following way

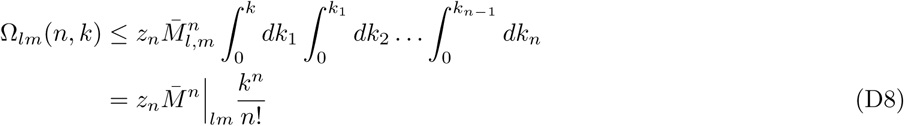

The matrix

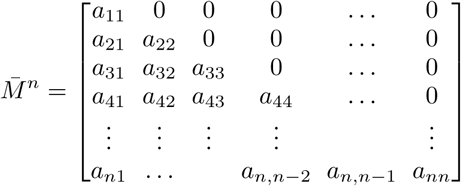

where *a*_*lm*_ = *S*_*lm*_ (*n*)*Ā*^*n*^ for appropriately defined polynomials *S*_*l,m*_ (*n*). Thus, it follows that 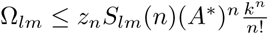 and 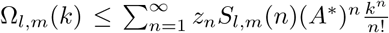. Consequently, the series will converge if *Āk* < 1, i.e. 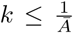. Assuming *µ*^(*j*)^ = *µ* ∀*j*, we can bound *Ā* as

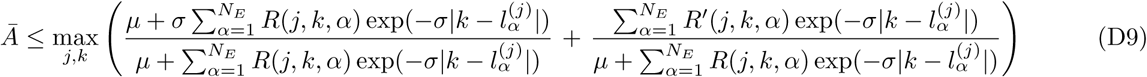

Since the parameters 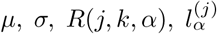 and *N*_*E*_ are finite and positive, and *R*′(*j, k, α*) is finite, *Ā* has a finite upper bound, implying *k* is always greater than zero, and the series has finite radius of convergence.

While in principle we can obtain the glycan profile for any *N*_*E*_ and *N*_*C*_ with arbitrary accuracy, assuming 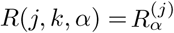, we provide explicit formulae for a few representative cases : (i) (*N*_*E*_ = 1, *N*_*C*_ = 1) and (ii) (*N*_*E*_ = 1, *N*_*C*_ = 2).

i. *N*_*E*_ = 1, *N*_*C*_ = 1: The solution of the differential equation is given by

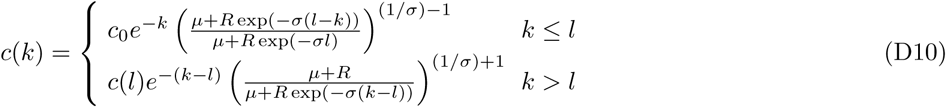 A representative concentration profile is plotted in Fig. A2(a). The concentration profile consists of two distinct components: an initial exponential decay, and then an exponential rise and fall concentrated around *l*. The relative weight of these two components is controlled by the sensitivity *σ* and the rate *R*. Such explicit formulae can be obtained for any *N*_*E*_ > 1, as long as *N*_*C*_ = 1.
ii. *N*_*E*_ = 1, *N*_*C*_ = 2: The concentration profile *c*^(2)^ in cisterna 2 can be obtained from the following calculation. Let *l*^(*j*)^ denote the “length” of the enzyme in cisterna *j* = 1, 2. For *k* ≤ min{*l*^(1)^, *l*^(2)^}

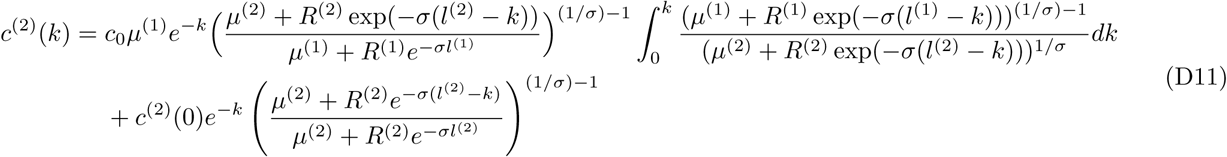

Next, consider the case where *l*^(1)^ ≤ *l*^(2)^. Then, for *l*^(1)^ < *k* ≤ *l*^(2)^

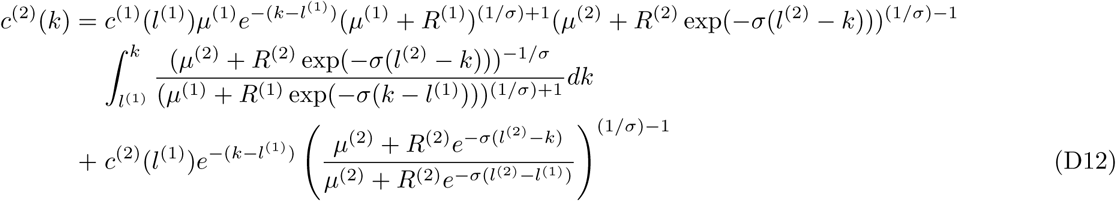

**FIG. A2.**
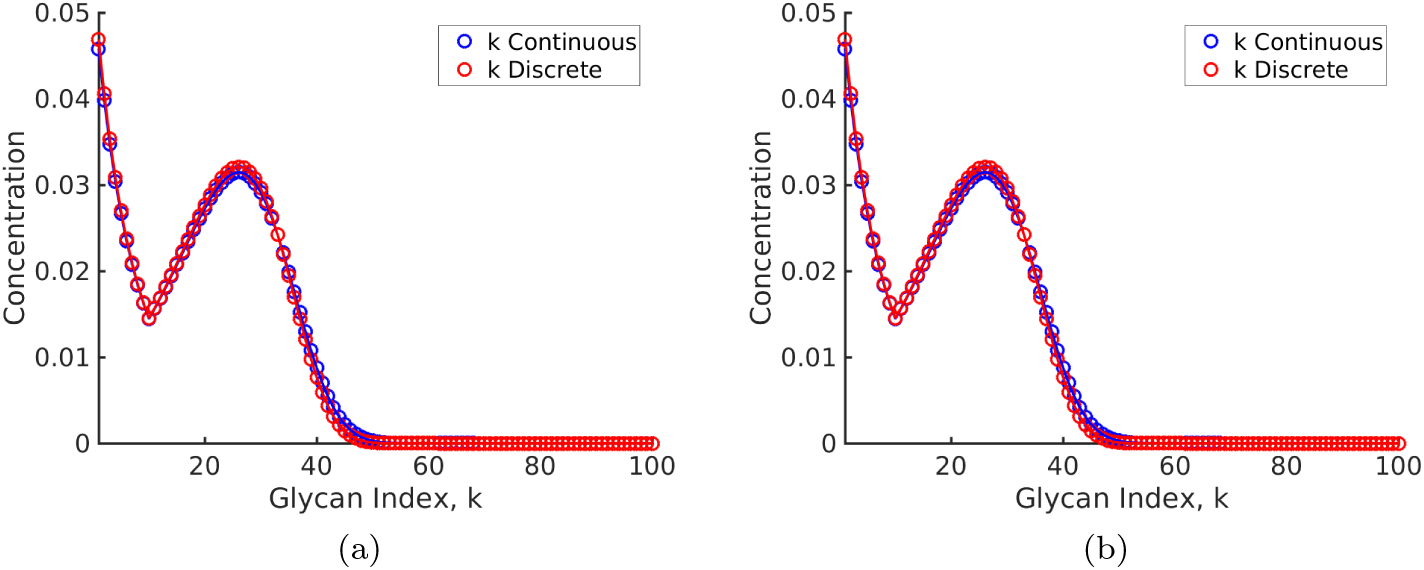
Glycan concentration profile calculated from the model using (a) formula (D10) for *N*_*E*_ = *N*_*C*_ = 1 and (b) formulae (D11)-(D15) for *N*_*E*_ = 1, *N*_*C*_ = 2.

and for *l*^(1)^ ≤ *l*^(2)^ < *k*,

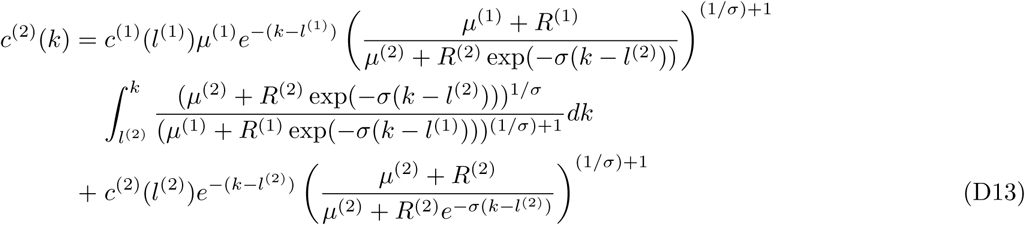

Next, the case where *l*^(1)^ ≥ *l*^(2)^. For *l*^(2)^ < *k* ≤ *l*^(1)^,

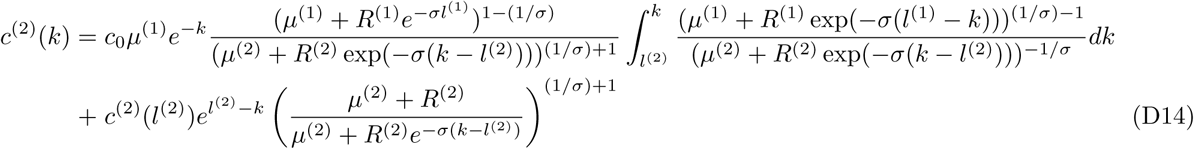

For ^(2)^ ≤ *l*^(1)^ < *k*,

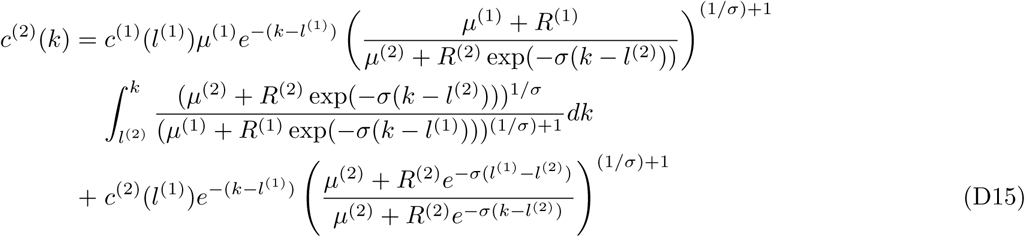

The integrals in (D11) to (D15) can evaluated numerically. The result of the numerical computation is shown in Fig. A2.

## Appendix E: Capability of the chemical network model to generate complex distributions

Is our glycan synthesis model capable of generating concentration distributions of arbitrary complexity? In what way do we need to change the parameters *N*_*E*_, *N*_*C*_, *σ*, …, in order to generate glycan distributions **c** of a given complexity?

The purpose of this section, is to obtain some heuristics for this task.

We show in Appendix D that (B4) can be solved analytically in the limit *N*_*s*_ ≫ 1, because in this limit the glycan index *k* can be approximated by a continuous variable, and the recursion relations for the steady state glycan concentrations (5)-(6) can be cast as a matrix differential equation. This allows us to obtain an *explicit* expression for the steady state concentration in terms of the parameters (**R, L**).

We derive our heuristics from a semi-analytical treatment in the limit *N*_*s*_ ≫ 1 (Appendix D), which apart from being simple to implement in general, provides an explicit formula for *c*_*k*_ for the case *N*_*E*_ = *N*_*C*_ = 1 (D10). Figures A3(a)-(d) show the glycan profile *c*_*k*_ *vs. k* as one varies the enzyme specificity *σ*, the reaction rates *R* and transport rates *µ*, for two different values of *N*_*E*_ and *N*_*C*_. The results in the plots lead us to the following general observations:

- Very low specificity enzymes cannot generate complex glycan distributions. Keeping everything else fixed, intermediate or high specificity enzymes can generate glycan distributions of higher complexity by increasing *N*_*E*_ or *N*_*C*_ (Figs. A3(a),(c)).
- Decreasing the specificity *σ* or increasing the rates *R* increases the proportion of higher index glycans. Keeping everything else fixed, changes in the rate *R* have a stronger impact on the relative weights of the higher index glycans to lower index glycans. The relative weight of the higher index glycans increases with increasing *N*_*E*_ and *N*_*C*_ (Figs. A3(b)-(d)).
- Keeping everything else fixed, decreasing enzyme specificity increases the spread of the distribution around the peaks (Figs. A3(a),(c)).

## Appendix F: Parameter estimation

The typical transport time of glycoproteins across the Golgi complex is estimated to be in the range 15-20 mins. [23], which corresponds to the transport rate, *µ* = .18/min. We bound the transport rate for our optimization between 0.01/min and 1/min.

Next, we estimate the range of values for the chemical reaction rates. The injection rate *q* is in the range 100 − 1500 pmol/10^6^ cell 24 h [23, 24]. For our calculation we set *q* = 387.30 pmol/10^6^ cells 24 hr = 0.27 pmol/10^6^ cells min, the geometric mean of 100 and 1500. We set the range for the enzymatic rate *R* to be

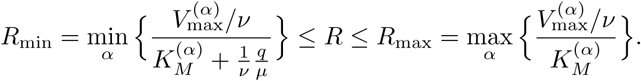

where 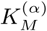 and 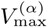 denote the Michaelis constants and *V*_max_ of the *α*-th enzyme. The conversion from 1 pmoles/10^6^ cells to concentration can be obtained by taking cisternal volume (*ν*) to be 2.5*µm*^3^ [23, 24]. This gives

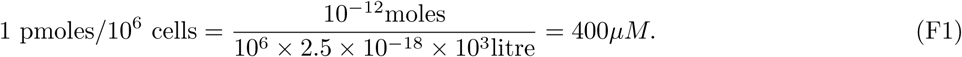

In Table I we report the parameters for the 8 enzymes taken from Table 3 in [23]. From these parameters it follows that

**TABLE I.**
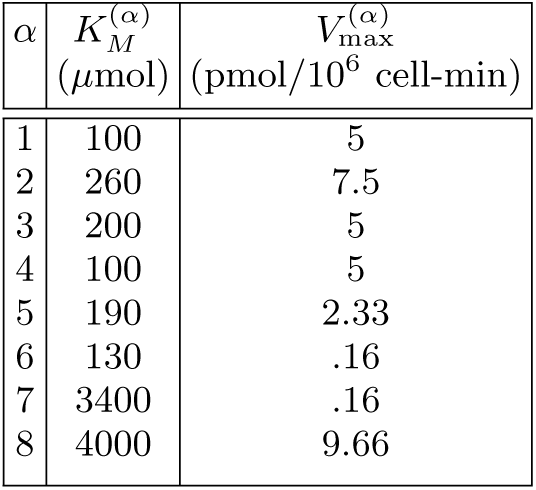
Enzyme parameters taken from Table 3 in [23] that we use to calculate the bounds on the reaction rate *R*. Here 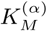 and 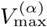 denote the Michaelis constant and *V* of the *α*-th enzyme.

| $\alpha$ | $K_M^{(\alpha)}$<br>($\mu\text{mol}$) | $V_{\max}^{(\alpha)}$<br>($\text{pmol}/10^6 \text{ cell-min}$) |
| --- | --- | --- |
| 1 | 100 | 5 |
| 2 | 260 | 7.5 |
| 3 | 200 | 5 |
| 4 | 100 | 5 |
| 5 | 190 | 2.33 |
| 6 | 130 | .16 |
| 7 | 3400 | .16 |
| 8 | 4000 | 9.66 |

**FIG. A3.**
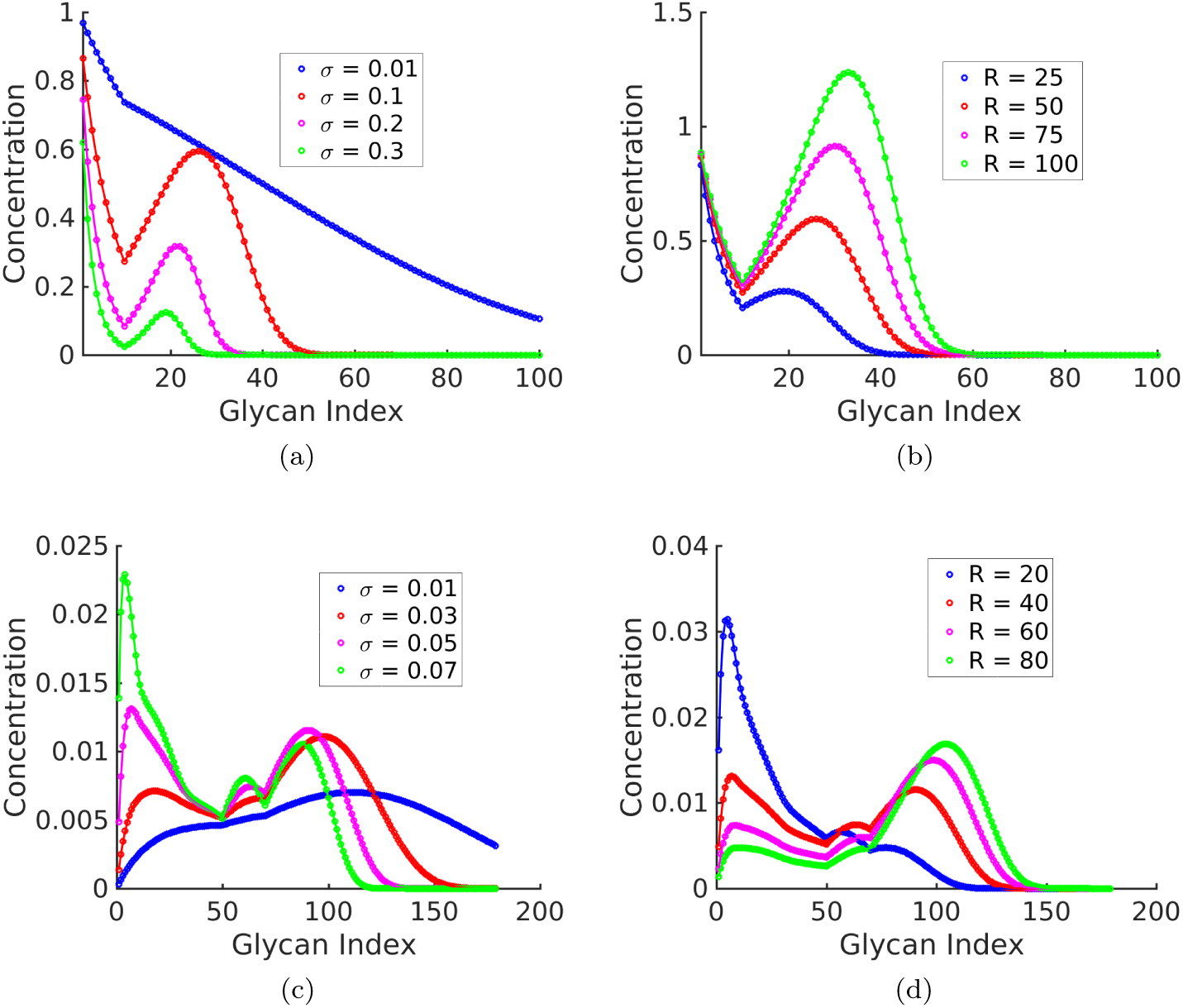
Glycan profile {*c*_*k*_ : *k* = 1, …, *N*_*s*_ } as a function of specificity *σ* (Fig. (a), (c)), and reaction rates *R* (Fig. (b), (d)). Fig. (a): *N*_*E*_ = *N*_*C*_ = 1, (*R* = 50, *µ* = 1, *l* = 10). *c*_*k*_ decreases exponentially with *k* for very low and very high *σ*; however, the decay rate is lower at low *σ*. For intermediate values of *σ*, the distribution has *exactly* two peaks, one of which is at *k* = 0, and eventually decays exponentially. The width of the distribution is a decreasing function of *σ*. Fig. (b): *N*_*E*_ = *N*_*C*_ = 1, (*σ* = 0.1, *µ* = 1, *l* = 10). At low *R, c*_*k*_ is concentrated at low *k*. The proportion of higher index glycans in an increasing function of *R*. Fig. (c): 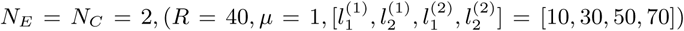. As *σ* increases, the distribution becomes more complex – from a single peaked distribution at low *σ* to a maximum of four-peaked distribution at high *σ*. The peaks gets sharper, and more well defined as *σ* increases. Fig. (d): 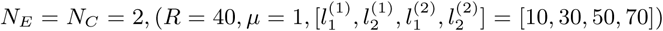. As in the plots in Fig. (b), increasing *R* shifts the peaks towards higher index glycans and the proportion of higher index glycan increases.

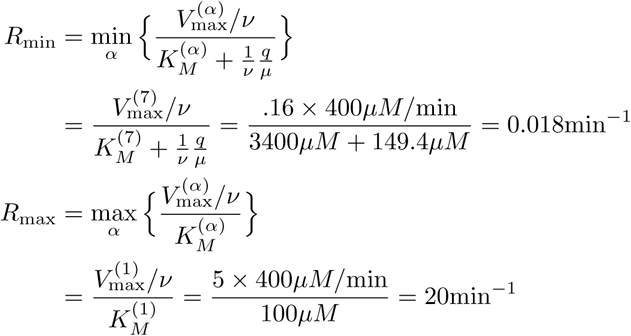

## Appendix G: Constructing target distributions for glycans of a given cell type

The target distribution of the glycans on the cell surface is obtained via mass spectrometry. The x-axis of mass spectroscopy (MS) graphs is mass/charge of the ionised sample molecules and the y-axis is relative intensity corresponding to each mass/charge value, taking the highest intensity as 100%.

This relative intensity roughly correlates with the relative abundances of the molecules in the sample.

This raw MS data is noisy and cannot be directly used as the target distribution in our optimization problem. There are three major sources of noise in the MS data [47]: the chemical noise in the sample, the Poisson noise associated with detecting discrete events, and the Nyquist-Johnson noise associated with any charge system. We propose a simple model that accounts for the chemical noise and the Poisson sampling noise. Using this noise model and the available MS data, we generate parametric bootstrap samples of glycan measurements, and fit a Gaussian Mixture Model (GMM) on this sample to approximate the glycan distribution. This GMM probability distribution is used as the target distribution in our numerical experiments.

The MS data obtained from [22] had mass ranging between 500 to 5000 Daltons with intensity reported at every 0.0153 Daltons. We first bin this MS data into 180 bins and take the maximum value within each bin as the value of intensity for that bin. Fig. A4 shows the raw MS data and the binned distribution.

Let *Ī*_*k*_ represents the relative intensity of the *k*-th bin in the binned MS graph. We generate a sample population of glycans using the MS data in the following way:

1. Poisson sampling noise: The MS data does not have absolute count information. We assume an arbitrary maximum count *I*_max_, and define the intensity *I*_*k*_ = *I*_max_*Ī*_*k*_. The plots in Fig. A5(a) show that the results are not sensitive to the specific value of *I*_max_.
2. Chemical noise: The sample used for MS analysis also contains small amounts of molecules that are not glycans. These appear as the very small peaks in the MS data. We assume that the probability *p*_*k*_ that the peak at index *k* corresponds to a glycan is given by

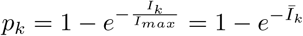

which adequately suppresses this chemical noise.
3. Bootstrapped glycan data: The count *n*_*k*_ at the glycan index *k* is distributed according to the following distribution:

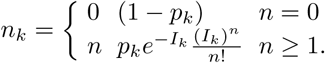

We assume that the MS data was generated from *N* different cells. Thus, the total count at glycan index *k* is given by the sum of *N*_*i*_ i.i.d. samples distributed according to the distribution above. We in Fig. A5 (b) show that results are insensitive to *N*_*i*_.

**FIG. A4.**
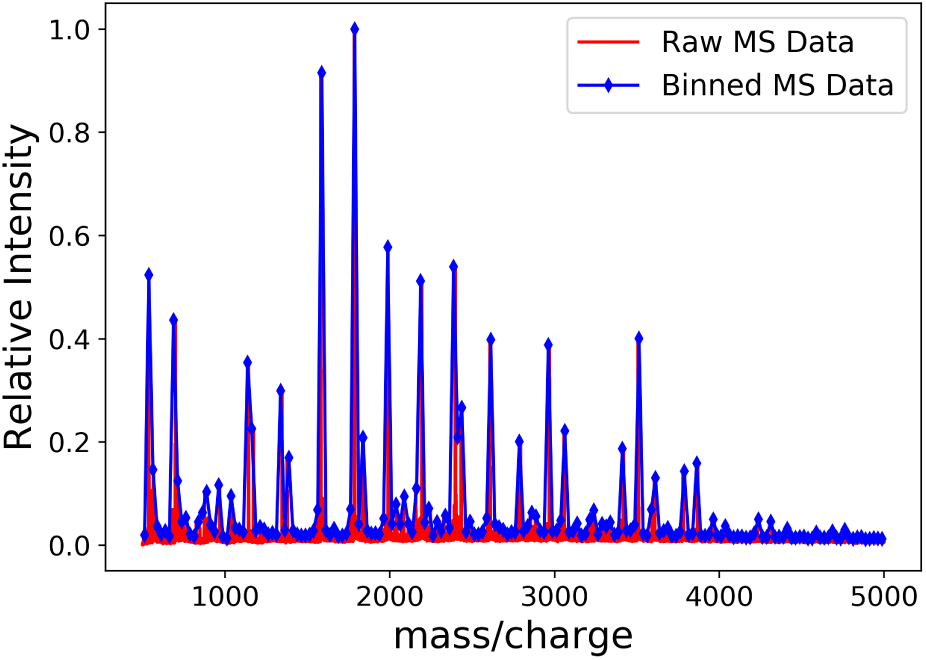
The binned MS data (blue) approximates the raw MS data (red) very well. We use this binned data for GMM approximation of the MS data.

**FIG. A5.**
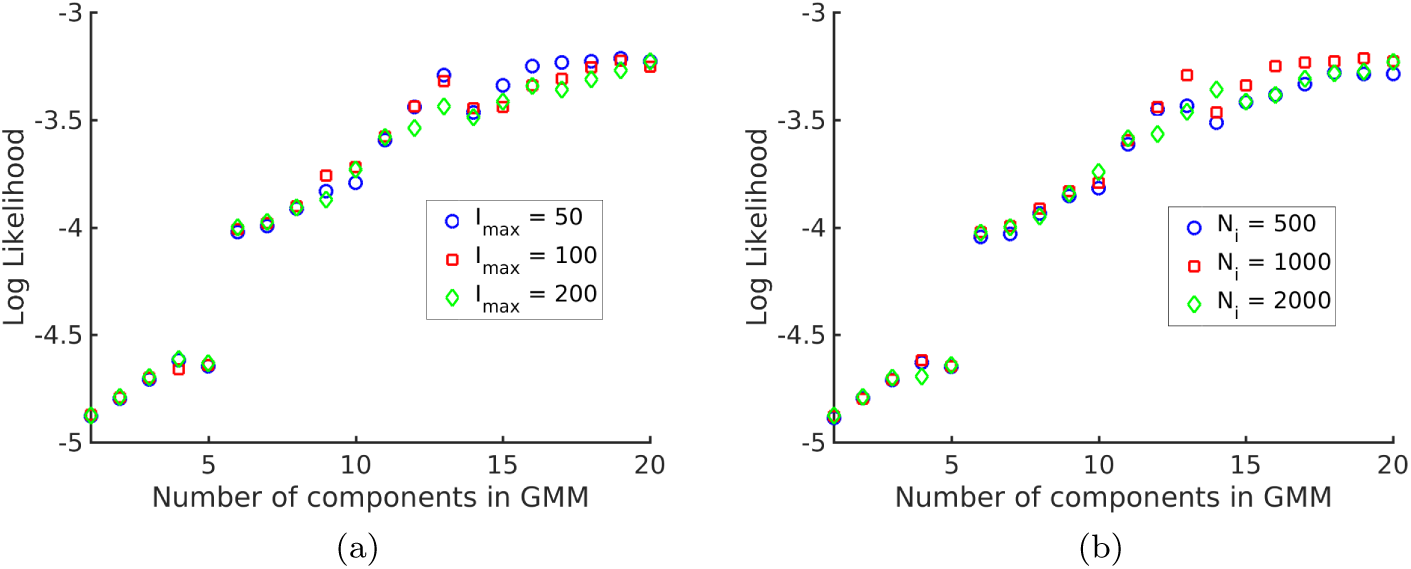
Log likelihood vs. number of components (*N*) in the GMM. We see that the log likelihood saturates at around *N* = 20, thus 20-GMM is a very good representation of the MS-data from *human* T-cells. The different symbols are for (a) different values of the maximum intensity *I*_*max*_ = 50, 100, 200 and (b) different values of the number of i.i.d samples *N*_*i*_ = 500, 1000, 2000, showing the insensitivity of the log likelihood to the value of *I*_*max*_ and *N*_*i*_.

Next, we interpret the counts as samples from a “spatial” distribution *f*. We approximate this distribution as a Gaussian mixture, i.e. 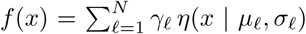, where *η*(*x* | *µ, σ*) denotes the density of a normally distributed random variable with mean *µ* and standard deviation *σ* at the location *x*. In this setting, we assume that each count is a sample from the distribution *η*(*x* | *µ*_*ℓ*_, *σ*_*ℓ*_) with probability *γ*_*ℓ*_. Thus, each count is classified as coming from one of the Gaussian components.

## Appendix H: Numerical scheme for performing the non-convex optimization

We solve Optimization C using the numerical scheme detailed below. The optimization problem consists of minimising a non-convex objective with linear box constraints. We use the MATLAB FMINCON function to solve this optimization. We use Sequential Quadratic Programming (SQP), a gradient based iterative optimization scheme for solving optimizations with non-linear differentiable objective and constraints. Since our problem is non-convex and SQP only gives local minima, we initialise the algorithm with many random initial points. We use SOBOLSET function of MATLAB to generate space filling pseudo random numbers. We have taken 1000 initialisations for each *N*_*E*_, *N*_*C*_ and *σ* value. We have taken 50 equally spaced points between 0 and 1 to explore the *σ*-space for Fig. 3. Some minor fluctuations in *D* due to non-convexity of the objective function in the final results were smoothed out by taking the convex hull of the *D* vs. *σ* graph. The results for *σ*_*min*_(*N*_*E*_, *N*_*C*_) and *D*(*σ*_*min*_, *N*_*E*_, *N*_*C*_) (Fig. 4) were obtained by adding *σ* to the optimization vector and then performing the optimization. The sensitivity results (Figs. 4e and 4e) were obtained by approximating the *D* vs *σ* graph around *σ*_*min*_ with a parabola, the coefficient of the quadratic term being the curvature of the graph at *σ*_*min*_.

**FIG. A6.**
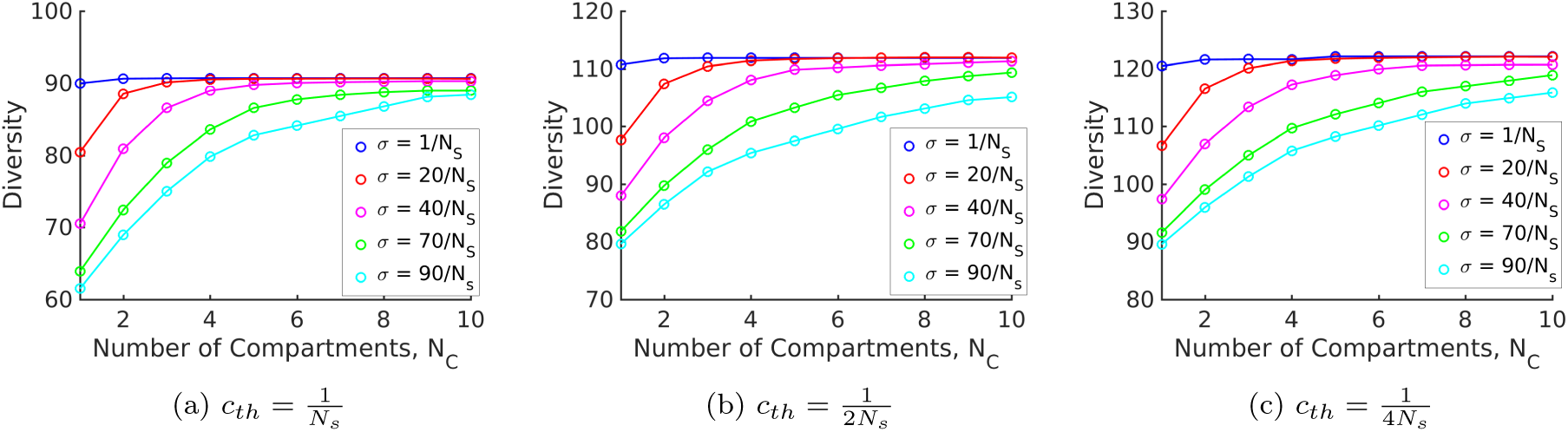
Diversity vs. *N*_*C*_ for different values of *σ* keeping *N*_*E*_ = 1 fixed, for three different values of the threshold, 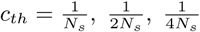. Changing the value of the threshold *c*_*th*_, only changes the saturation value of the diversity curve.

A similar numerical scheme was used to optimize diversity.

## Footnotes

1 A rigorous definition of complexity can be given in terms of the Kullback-Leibler metric [16, 17] between two glycan profiles. We declare that two profiles are distinguishable only if the Kullback-Leibler distance between the profiles is more than a given tolerance. This tolerance is an increasing function of the noise. We define the *complexity* of a set of possible glycan profiles as the size of the largest subset such that the Kullback-Leibler distance of any pair of profiles is larger than the tolerance.

2 We normalize the distribution so that 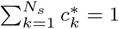

3 In Appendix D we show that (B4) can be solved analytically in the limit *N*_*s*_ ≫ 1, since the glycan index *k* can be approximated by a continuous variable, and the recursion relations for the steady state glycan concentrations (5)-(6) can be cast as a matrix differential equation. This allows us to obtain an *explicit* expression for the steady state concentration in terms of the parameters (**R, L**).

